# Data-based RNA-seq Simulations by Binomial Thinning

**DOI:** 10.1101/758524

**Authors:** David Gerard

**Affiliations:** Department of Mathematics and Statistics, American University, Washington, DC, 20016, USA

**Keywords:** RNA-seq, simulation, differential expression, factor analysis, confounders, scaling factors

## Abstract

With the explosion in the number of methods designed to analyze bulk and single-cell RNA-seq data, there is a growing need for approaches that assess and compare these methods. The usual technique is to compare methods on data simulated according to some theoretical model. However, as real data often exhibit violations from theoretical models, this can result in unsubstantiated claims of a method’s performance. Rather than generate data from a theoretical model, in this paper we develop methods to add signal to real RNA-seq datasets. Since the resulting simulated data are not generated from an unrealistic theoretical model, they exhibit realistic (annoying) attributes of real data. This lets RNA-seq methods developers assess their procedures in non-ideal (model-violating) scenarios. Our procedures may be applied to both single-cell and bulk RNA-seq. We show that our simulation method results in more realistic datasets and can alter the conclusions of a differential expression analysis study. We also demonstrate our approach by comparing various factor analysis techniques on RNA-seq datasets. Our tools are available in the `seqgendiff` R package on the Comprehensive R Archive Net-work: https://cran.r-project.org/package=seqgendiff.

## 1 Introduction

Due to its higher signal-to-noise ratio, larger range of detection, and its ability to measure *a priori* unknown genes, RNA-seq has surpassed microarrays as the technology of choice to measure gene expression [Wang et al., 2009]. With the advent of single-cell RNA-seq technologies, researchers now even have the ability to explore expression variation at the individual cell level [Hwang et al., 2018]. This presents exciting opportunities for researchers to characterize the expression heterogeneity between and within organisms, and has brought about a plentiful flow of new datasets. In the wake of these new data, an explosion of methods has been developed to analyze them. In Sections 2.2, 2.3, 2.4, and 2.5 we provide a large (yet terribly incomplete) list of methods designed to analyze RNA-seq data.

The typical pipeline to evaluate a method is to first simulate data according to some theoretical model, then compare it to competing methods on these simulated data and show it to be superior in some fashion. This way of evaluation can be useful to see how a method works in ideal scenarios. However, real data rarely live in ideal scenarios. Real data often exhibit unwanted variation beyond that assumed by a model [Leek et al., 2010]. Theoretical distributional assumptions are also difficult to verify, and are sometimes mired in controversy [Svensson, 2019].

In this paper, we propose an alternative approach. Rather than generate data with a prespecified signal according to some modeling assumptions, we take a real RNA-seq dataset and add a prespecified signal to it. The main advantage of our approach is that any unwanted variation in the real data is maintained in the simulated data, and this unwanted variation need not be prespecified by the researcher. The way we add signal does carry assumptions, but they are flexible (Supplementary Section S1.2). And we believe that this way of simulation, compared to simulating under a theoretical model, allows researchers to more realistically evaluate their methods.

This manuscript essentially generalizes the simulation techniques proposed in Gerard and Stephens [2017], Gerard and Stephens [2018], and Lu [2018]. These previous papers use binomial thinning (the approach used in this paper) in the case where there are just two groups that are differentially expressed (hereafter, the “two-group model”). These papers did not develop methods for more complicated design scenarios, they did not present user-friendly software implementations for their simulation techniques, and they did not justify their simulation techniques as broadly. Here, we allow for arbitrary experimental designs, we release software for users to implement their own simulations, and we justify our techniques using very flexible assumptions.

There has been some other previous work on “data-based” simulations in expression analyses. Datasets resulting from data-based simulations (sometimes called “plasmodes” [Mehta et al., 2004]) have been used in microarray studies before the development of RNA-seq [Nettleton et al., 2007, Gadbury et al., 2008]. All RNA-seq data-based simulation methods have so far operated in the two-group (or finite-group) model, without any ability to simulate data from arbitrary experimental designs. Rocke et al. [2015] and Sun and Stephens [2018] randomly shuffled group indicators in the two-group model, resulting in completely null data, and methods can be evaluated on their ability to control for type I error when the data are all null. Rigaill et al. [2016], in addition to generating null data by randomly shuffling group labels, incorporate multiple datasets to create some non-null genes within their simulated datasets. Benidt and Nettleton [2015] use a count-swapping algorithm in the two-group model to create differentially expressed genes when one already has two treatment groups. Kvam et al. [2012], Reeb and Steibel [2013], and van de Wiel et al. [2014] create non-null genes by multiplying counts for all individuals in a group by the fold-change in mean expression. Robinson and Storey [2014] uses a binomial distribution approach to uniformly decrease the sequencing depth of an entire dataset (but not to add differentially expressed genes). Concerning non-data-based methods, Vieth et al. [2017] and Zappia et al. [2017] use real RNA-seq data to estimate the parameters in a data-generating model before simulating data from the theoretical model using these estimated parameter values. Our work is the first to extend data-based RNA-seq simulation beyond the finite-group model.

Our paper is organized as follows. We first list the goals and assumptions of our simulation scheme (Section 2.1) before motivating it with four applications (Sections 2.2, 2.3, 2.4, and 2.5) and describing our process of simulating RNA-seq in detail (Section 2.6). We then demonstrate how our approach can more accurately preserve structure in a real dataset compared to simulating a dataset from a theoretical model (Section 3.1). We show that this can alter the conclusions of a differential expression analysis simulation study (Section 3.2). We then apply our simulation approach by comparing five factor analysis methods using the GTEx data [GTEx Consortium, 2017] (Section 3.3). We finish with a discussion and conclusions (Sections 4 and 5).

We adopt the following notation. We denote matrices by bold uppercase letters (***A***), vectors by bold lowercase letters (***a***), and scalars by non-bold letters (*a* or *A*). Indices typically run from 1 to their uppercase version, e.g. *a* = 1, 2, …, *A*. Where there is no chance for confusion, we let non-bold versions of letters represent the scalar elements of matrices and vectors. So *a*_*ij*_ is the (*i, j*)th element of ***A***, while *a*_*i*_ is the *i*th element of ***a***. We let **1**_*A*_ denote the *A*-vector of 1’s and **1**_*A*×*B*_ the *A* × *B* matrix of 1’s. The matrix transpose is denoted by ***A***^T^.

## 2 Methods

### 2.1 Goals and Assumptions

We will now describe the goals and assumptions of our simulation method, which relies on a researcher having access to a real RNA-seq dataset. Suppose a researcher has a matrix ***Y*** ∈ ℝ^*G*×*N*^ of RNA-seq read-counts for *G* genes and *N* individuals. Also suppose a researcher has access to a design matrix 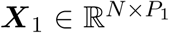 with *P*_1_ variables. We assume the RNA-seq counts, ***Y***, are generated according to the following model:

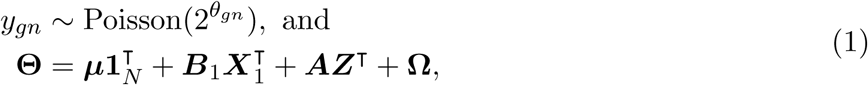

where

- ***µ*** ∈ ℝ^*G*^ is a vector of intercept terms for the genes,
- 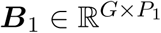 is the corresponding coefficient matrix of ***X***_1_,
- ***Z*** ∈ ℝ^*N*×*K*^ is a matrix of unobserved surrogate variables,
- ***A*** ∈ ℝ^*G*×*K*^ is the corresponding coefficient matrix of ***Z***, and
- **Ω** ∈ ℝ^*G*×*N*^ represents all other unwanted variation not accommodated by the other terms in the model,

where ***µ***, ***B***_1_, ***Z***, ***A***, and **Ω** are all unknown. Given the above data-generating process, suppose a user provides the following (known) elements:

- 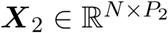, a design matrix with fixed-rows (see note 3 below),
- 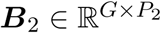, the coefficient matrix corresponding to ***X***_2_,
- 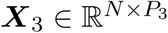, a design matrix with rows that can be permuted (see note 3 below), and
- 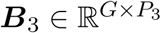, the coefficient matrix corresponding to ***X***_3_.

Our goal is to generate a matrix 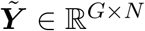 from ***Y*** such that

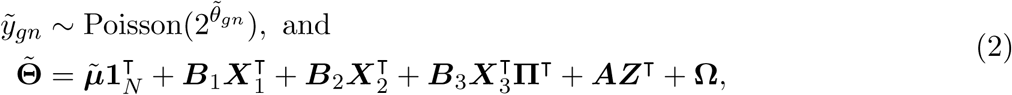

where

- **Π** ∈ ℝ^*N*×*N*^ is a random permutation matrix, whose distribution controls the level of association between the columns of **Π*X***_3_ and the columns of ***Z***, and
- 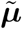 is a new vector of intercept terms for the genes.

We will provide the details on how to generate 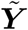 from ***Y*** in Section 2.6. But we would like to first provide some notes below, and then discuss the applications of being able to generate (2) from (1).

**Note 1:** For simplicity we use the Poisson distribution in the main text (equations (1) and (2)). However, our approach is valid under much more general assumptions. In particular, we note that if the counts were generated according to a negative binomial distribution, a zero-inflated negative binomial distribution, or even a mixture of binomials and negative binomials, then our simulation scheme still preserves the structure of the data (Supplementary Section S1.2). However, even when our general modeling assumptions are violated, one can show (via the law of total expectation) that if log_2_(*E*[***Y***]) = **Θ**, then 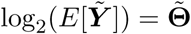, where we are taking element-wise logarithms of E[***Y***] and 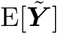. Thus, our procedure will produce the correct mean log_2_-fold change in the new dataset, but the resulting mean/variance relationship might not be as assumed.

**Note 2:** The **Ω** term in (1) and (2) represents the realistic and annoying features of the data. In ideal situations, **Ω** = **0**_*G*×*N*_. However, most datasets likely include non-zero **Ω**, and so assessing a method’s ability to be robust in the presence of **Ω**, without the researcher having to prespecify **Ω**, is the key strength of our simulation approach.

**Note 3:** As described below, we include both ***X***_2_ and ***X***_3_ in (2) to control different aspects of a simulation study. One may control the level of association between the columns of ***X***_1_ and ***X***_2_ as these are both observed and fixed by the user. The inclusion of ***X***_3_ and **Π** allows us to try to control the level of association between **Π*X***_3_ and ***Z***.

Before we discuss obtaining (2) from (1), we point out four potential applications of this simulation approach: (i) evaluating differential expression analyses (Section 2.2), (ii) evaluating confounder adjustment approaches (Section 2.3), (iii) evaluating the effects of library size heterogeneity on differential expression analyses (Section 2.4), and (iv) evaluating factor analysis methods (Section 2.5).

### 2.2 Application: Evaluating Differential Expression Analysis

One of the more common applications of RNA-seq data is estimating and testing for differences in gene expression between two groups. Many packages and techniques exist to perform this task [Robinson and Smyth, 2007b, Hardcastle and Kelly, 2010, Van De Wiel et al., 2012, Kharchenko et al., 2014, Law et al., 2014, Love et al., 2014, Finak et al., 2015, Guo et al., 2015, Nabavi et al., 2015, Delmans and Hemberg, 2016, Korthauer et al., 2016, Costa-Silva et al., 2017, Qiu et al., 2017, Miao et al., 2018, Risso et al., 2018, Van den Berge et al., 2018, Wang and Nabavi, 2018, Wang et al., 2019, among others], and so developing approaches and software to compare these different software packages would be of great utility to the scientific community. Generating data from the two-group model is a special case of (1) and (2), where

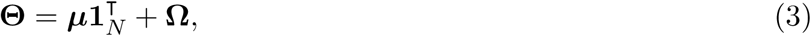

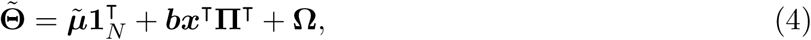

and **Π*x*** ∈ ℝ^*N*^ contains a single indicator variable, indicating membership to one of two groups. Researchers may specify ***b*** and ***x*** and evaluate a method’s ability to (i) estimate ***b*** and (ii) detect which genes have non-zero *b*_*g*_.

In many settings, a researcher may want to specify the distribution of the *b*_*g*_’s. Our software implementation allows for this. In addition, following Stephens [2016], we allow researchers to specify the distribution of 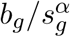, where *s*_*g*_ is the sample standard deviation of the *g*th row of log_2_(***Y*** + 0.5), and *α* is a user-specified constant. Allowing for *α* = 0 corresponds to the scenario of specifying the distribution of the effects, while allowing for *α* = 1 corresponds to specifying the *p*-value prior of Wakefield [2009].

Though the two-group model is perhaps the most common scenario in differential expression analysis, our method also allows for arbitrary design matrices. Such design matrices have applications in many types of expression experiments [Smyth, 2004, McCarthy et al., 2012, Van De Wiel et al., 2012, Tang et al., 2015], and so the ability to simulate arbitrary designs gives researchers another tool to evaluate their methods in more complicated scenarios.

### 2.3 Application: Evaluating Confounder Adjustment

Unobserved confounding / batch effects / surrogate variables / unwanted variation has been recognized as a serious impediment to scientific studies in the modern “omics” era [Leek et al., 2010]. As such, there is a large literature on accounting for unwanted variation, particularly in RNA-seq studies [Leek and Storey, 2007, Carvalho et al., 2008, Kang et al., 2008a,b, Leek and Storey, 2008, Stegle et al., 2008, Friguet et al., 2009, Kang et al., 2010, Listgarten et al., 2010, Stegle et al., 2010, Wu and Aryee, 2010, Fusi et al., 2012, Gagnon-Bartsch and Speed, 2012, Stegle et al., 2012, Sun et al., 2012, Gagnon-Bartsch et al., 2013, Mostafavi et al., 2013, Yang et al., 2013, Leek, 2014, Risso et al., 2014, Perry and Pillai, 2015, Chen and Zhou, 2017, Gerard and Stephens, 2017, Lee et al., 2017, Wang et al., 2017, Caye et al., 2018, Gerard and Stephens, 2018, Hung, 2018, McKennan and Nicolae, 2018a,b, among others]. The glut of available methods indicates a need to realistically compare these methods.

Typically, the form and strength of any unobserved confounding is not known. So one way to assess different confounder adjustment methods would be to assume model (1) and add signal to the data resulting in the following submodel of (2):

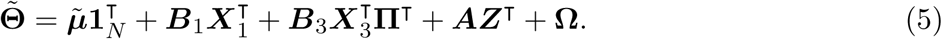

A researcher would then explore how close a method’s estimate of ***B***_3_ is to the truth (assuming the researcher may use both ***X***_1_ and **Π*X***_3_ to obtain this estimate). The researcher can control the correlation between the columns of **Π*X***_3_ and the columns of ***Z*** by specifying the distribution of **Π** (as described in Section 2.6). Intuitively, the stronger the correlation between the columns of ***X***_3_ and the columns of ***Z***, the more difficult the confounder adjustment problem. This approach was used in the two-group model in Gerard and Stephens [2017] and [2018], but not for general design matrices.

### 2.4 Application: Evaluating Effects of Library Size Heterogeneity

“Library size” corresponds to the number of reads an individual sample contains. Adjusting for library size is surprisingly subtle and difficult, and thus many techniques have been proposed to perform this adjustment [Anders and Huber, 2010, Bullard et al., 2010, Robinson and Oshlack, 2010, Langmead et al., 2010, Dillies et al., 2012]. The most commonly-used techniques can be viewed as a form of confounder adjustment [Gerard and Stephens, 2017]. For most methods, this form of confounder adjustment corresponds to setting one column of ***A*** in (1) to be **1**_*G*_ and estimating the corresponding column in ***Z*** using some robust method that assumes that the majority of genes are non-differentially expressed.

One way to evaluate the performance of a library size adjustment procedure is to see how effect size estimates change when the samples are thinned, changing the library size. First, assume we are operating in the following submodel of (1):

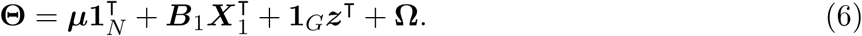

A researcher may specify (i) additional signal and (ii) a further amount of thinning on each sample by generating the following submodel of (2):

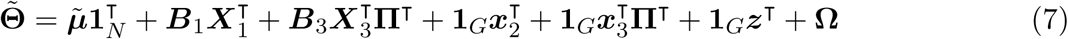

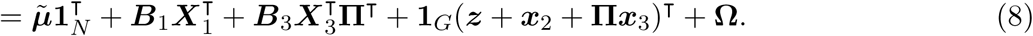

To evaluate the effectiveness of a library size adjustment procedure, researchers may observe the effects on the estimates of ***B***_3_ under various amounts of library thinning (controlled by altering ***x***_2_ and ***x***_**3**_).

### 2.5 Application: Evaluating Factor Analysis

Factor analysis is a fundamental technique in every statistician’s arsenal. Since its creation by Spearman [Spearman, 1904], literally hundreds of factor analysis / matrix decomposition / matrix factorization approaches have been developed, and new approaches are created each year to account for new features of new data [Hotelling, 1933, Eckart and Young, 1936, Comon, 1994, Tipping and Bishop, 1999, Lee and Seung, 1999, Hyvärinen and Oja, 2000, West, 2003, Zou et al., 2006, Hoff, 2007, Salakhutdinov and Mnih, 2008, Ghosh and Dunson, 2009, Witten et al., 2009, Engelhardt and Stephens, 2010, Stegle et al., 2010, Mayrink and Lucas, 2013, Yang et al., 2014, Josse and Wager, 2016, Leung and Drton, 2016, Wang and Stephens, 2018, to name a very few]. For RNA-seq, factor analysis methods have found applications in accounting for unwanted variation [Leek, 2014, Risso et al., 2014], estimating cell-cycle state [Buettner et al., 2015, Scialdone et al., 2015], and general quality assessments [Love et al., 2014]. Thus, techniques to realistically compare various factor analysis methods would be of great use to the scientific community. We demonstrate in this section how our simulation approaches can be used to evaluate factor analysis methods applied to RNA-seq.

We suppose that the RNA-seq read-counts follow the following submodel of (1):

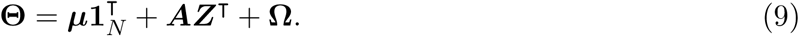

We then suppose that the researcher generates a modified dataset that follows the following sub-model of (2):

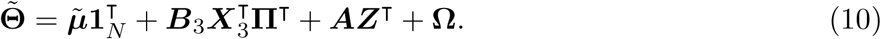

We assume that a researcher applies a factor analysis to (10) to estimate a low-rank matrix with *K* + *P*_3_ factors. That is, the researcher fits the following model,

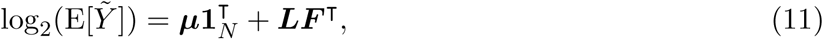

with factor matrix 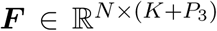 and loading matrix 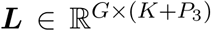, obtaining estimates 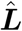 and 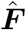. These estimates are obtained without using **Π*X***_3_. A researcher may evaluate their factor analysis by

1. Assessing if any of the columns of 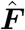 are close to the columns of **Π*X***_3_,
2. Assessing if any of the columns of 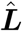 are close to the columns of ***B***_3_, and
3. Assessing if the column-space of **Π*X***_**3**_ is close to the column-space of 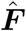, which would be an important consideration in downstream regression analyses [Leek and Storey, 2007, e.g.].

In a factor analysis, the factors and loadings are only identifiable after imposing assumptions on their structure (such as sparsity or orthogonality). Thus, researchers may vary the structure of ***B***_3_ and **Π*X***_3_ and observe the robustness of their factor analysis methods to violations of their structural assumptions.

### 2.6 Generating Modified RNA-seq Data

We will now discuss the approach of obtaining (2) from (1). We will use the following well-known fact of the Poisson distribution, which may be found in many elementary probability texts:

#### Lemma 1.

*If y ∼* Poisson(*a*) *and* 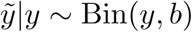, *then* 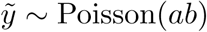.

In the case when **Π** is drawn uniformly from the space of permutation matrices, we have the simplified procedure described in Procedure 1. The validity of Procedure 1 follows directly from the modeling assumptions in (1) and Lemma 1. Since 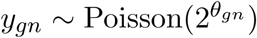 and 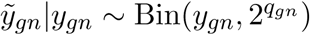, we have that 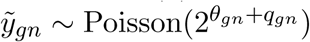. If we set 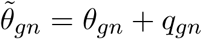, then we have

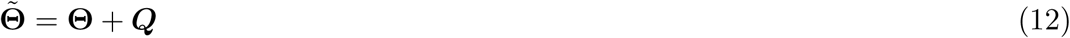

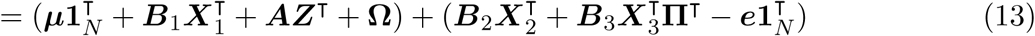

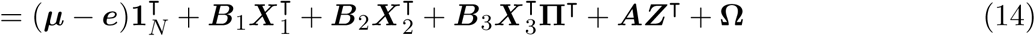

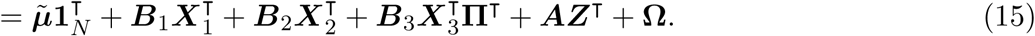

Equation (13) follows from the definition of **Θ** from (1) and the definition of ***Q*** from Step 4 of Procedure 1. Equation (15) follows by setting 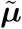 to be ***µ*** − ***e***.

#### Procedure 1

Basic procedure to generate (2) from (1) when the permuted design matrix (**Π*X***_3_) is independent of the surrogate variables.

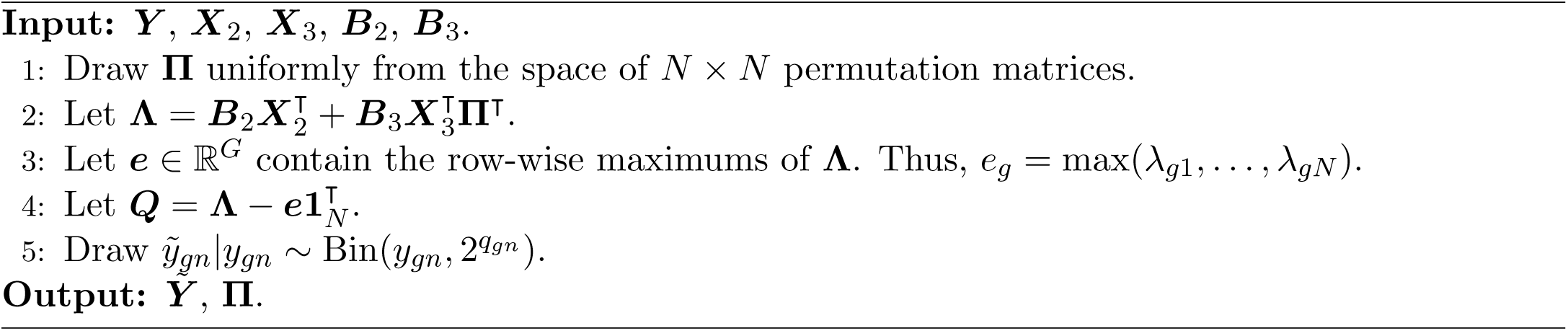

There are two main reasons to subtract the row-wise maximum from each row in Step 4 of Procedure 1: (i) this ensures that the binomial probabilities 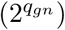 are always between 0 and 1, and (ii) this allows for minimal count-thinning while still obtaining our goal of (2). That is, the binomial probabilities will all be between 0 and 1, but they will be as close to 1 as possible while still yielding (2), thereby reducing the amount of discarded counts.

The main disadvantage to Procedure 1 is that the surrogate variables (***Z***) will be independent of the user-specified covariates (**Π*X***_3_). To allow the user to control the level of association between the surrogate variables and the user-provided variables, we propose using Procedure 2 to choose **Π**, rather than drawing **Π** uniformly from the space of permutation matrices. In brief, the user specifies a “target correlation” matrix, 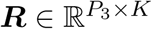, where *r*_*ik*_ is what the user desires to be the correlation between the *i*th column of **Π*X***_3_ and the *k*th column of ***Z***. We then estimate the surrogate variables either using a factor analysis (such as the truncated singular value decomposition) or surrogate variable analysis [Leek and Storey, 2007, 2008]. We then draw a new random matrix 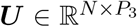 from a conditional normal distribution assuming that each row of ***U*** and ***Z*** is jointly normal with covariance matrix (16), thus the correlation between the columns of ***U*** and ***Z*** will be approximately ***R***. We then match the rows of ***X***_3_ with the rows of ***U*** using the pair-wise matching algorithm of Hansen and Klopfer [2006], though our software provides other options to match pairs via either the Gale-Shapley algorithm [Gale and Shapley, 1962] or the Hungarian algorithm [Kuhn, 1955]. This ensures that **Π*X***_3_ is as close to ***U*** as possible. We denote the permutation matrix that matches the rows of ***X***_3_ with the rows of ***U*** by **Π**.

#### Procedure 2

Procedure to draw a permutation matrix such that the surrogate variables are correlated with the permuted design matrix.

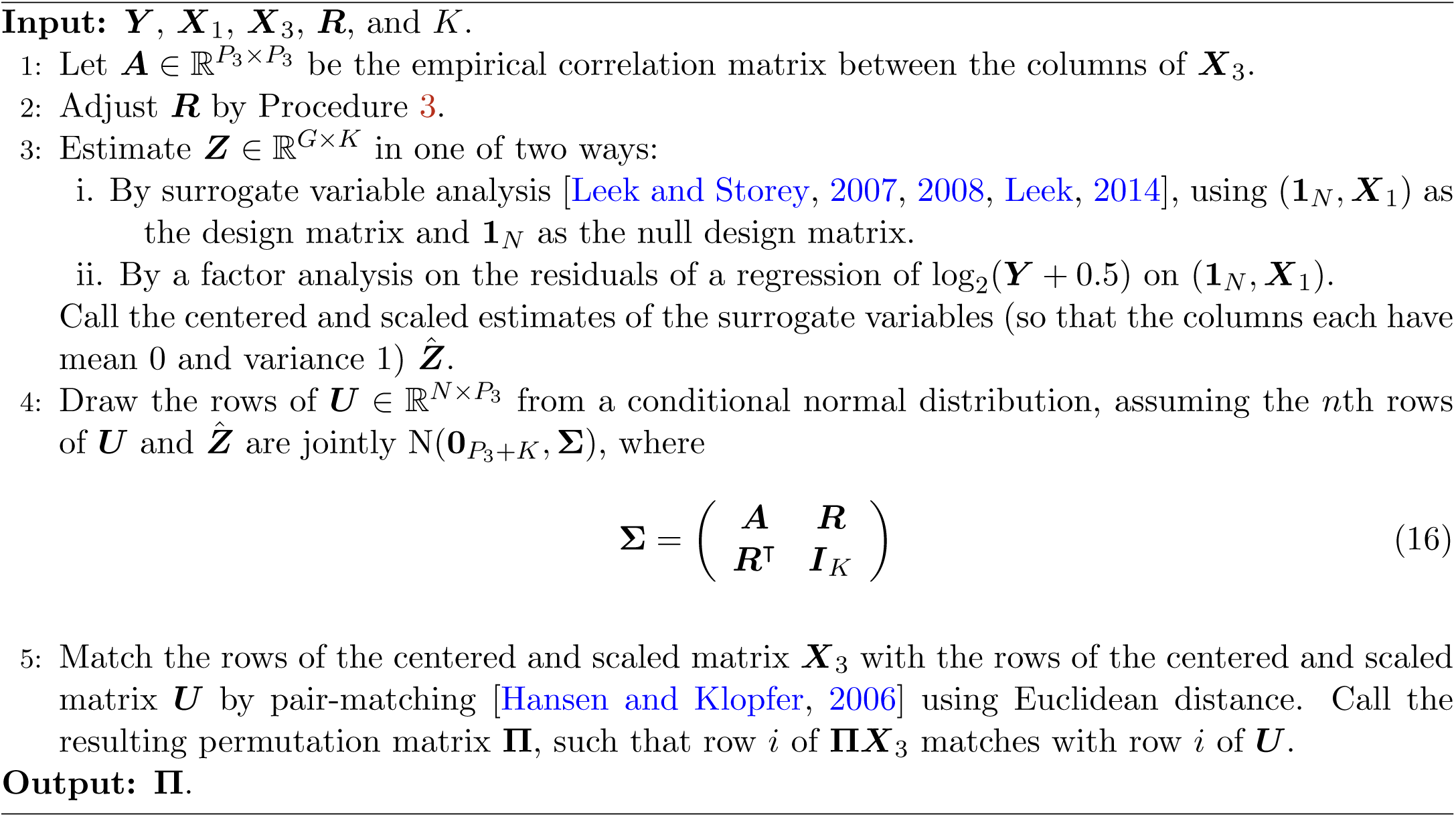

The resulting covariance matrix (16) used in Procedure 2 is not guaranteed to be positive semidefinite. Rather than demand the user specify an appropriate target correlation matrix (which might be in general difficult for the typical user), we modify the target correlation matrix using Procedure 3 to iteratively shrink ***R*** until the Schur complement condition for positive semi-definiteness [Zhang, 2006] is satisfied.

Procedure 2 is a compromise between letting the user specify the full design matrix ***X***_3_ and letting the user specify the correlation between the columns of **Π*X***_3_ and ***Z***. A user might want to specify the correlation between **Π*X***_3_ and ***Z*** to evaluate factor analyses in the presence of correlated factors (Section 2.5), or to evaluate how well confounder adjustment approaches cope in the presence of correlated confounders (Section 2.3). In the simple case when ***X***_3_ and 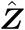 are drawn from a normal distribution, Procedure 2 will permute the rows of ***X***_3_ so that **Π*X***_3_ and 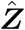 consistently has the correct correlation structure (Theorem 1). However, for general design matrices this will not be the case. Procedure 4 (implemented in our software) provides a Monte Carlo algorithm to estimate the true correlation given the target correlation. Basically, the estimator approximates the expected value (conditional on 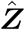) of the Pearson correlations between the columns of **Π*X***_3_ and the columns of 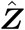. We justify this in an intuitive way by the law of total expectation. Consider ***x*** a single column of **Π*X***_3_ with empirical mean and standard deviation of 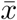 and *s*_*x*_. Similarly consider ***z*** a single column of 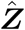 with empirical mean and standard deviation of 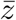 and *s*_*z*_. Then

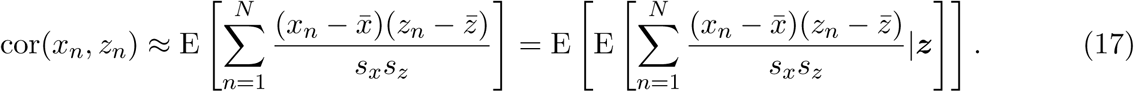

#### Procedure 3

Procedure to scale the target correlation matrix so that the overall correlation matrix is positive semi-definite.

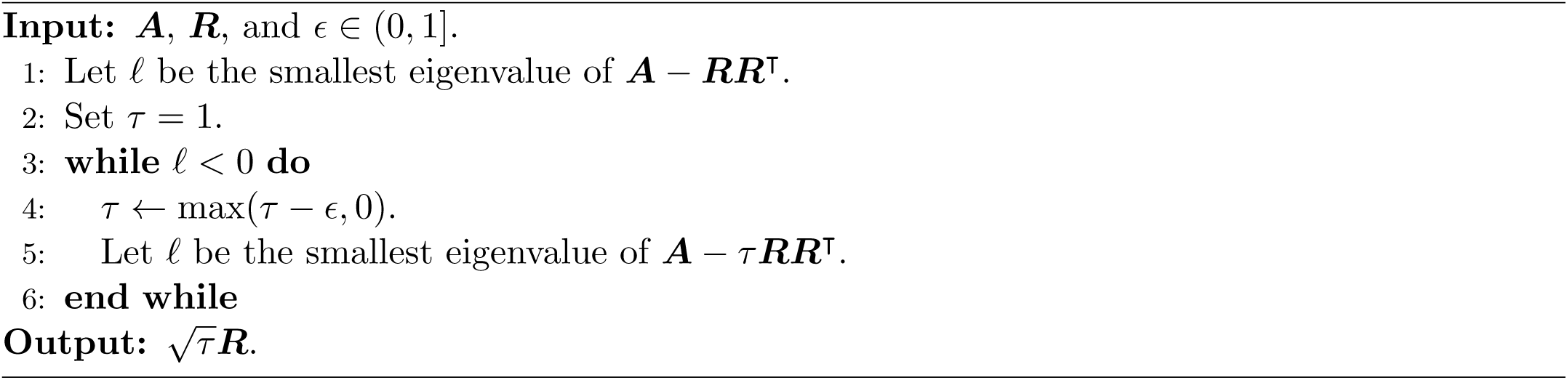

The estimator in Procedure 4 is a Monte Carlo approximation to the internal expectation in (17). We explore this correlation estimator through simulation in Supplementary Section S2.1.

#### Procedure 4

Monte Carlo procedure to estimate the true correlation matrix given the target correlation matrix.

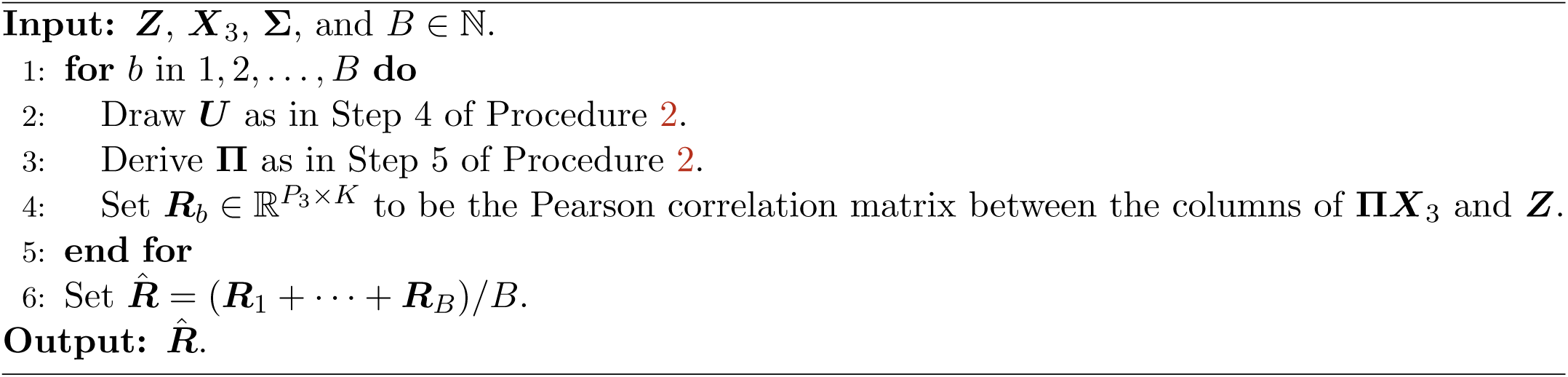

## 3 Results

### 3.1 Features of Real Data

Real data exhibit characteristics that are difficult to capture by simulations. In this section, we demonstrate how our binomial thinning approach maintains these features, while simulating from a theoretical model results in unrealistic simulated RNA-seq data.

We took the GTEx muscle data [GTEx Consortium, 2017], and filtered out all genes with a mean read-depth of less than 10 reads. This resulted in a dataset containing 18,204 genes and 564 individuals. We then randomly assigned half of the individuals to one group and half to the other group, and used our `seqgendiff` software to add a *N* (0, 0.8^2^) log_2_-fold-change between groups to 25% of the genes. We similarly used the `powsimR` software [Vieth et al., 2017] to generate data according to a theoretical negative binomial model (with parameters estimated from the GTEx muscle data), again by adding a *N* (0, 0.8^2^) log_2_-fold-change between the two groups in 25% of the genes. The results below are from one simulation, but the results are robust and consistent across many datasets. The reader is encouraged to change the random seed in our code to explore the robustness of our conclusions.

The structure of the `powsimR` dataset is very different from that observed in the `seqgendiff` and GTEx datasets. There seems to be more zeros in the `powsimR` dataset than in the `seqgendiff` and GTEx datasets (Supplementary Figure S2), even though we simulated the `powsimR` dataset under the negative binomial setting and not the zero-inflated negative binomial setting. Scree plots of the three datasets show that there are a lot more small factors influencing variation in the `seqgendiff` and GTEx datasets than in the `powsimR` dataset (Figure 1). The main source of variation in the `powsimR` dataset comes from the group membership, while other (unwanted) effects dominate the variation in the `seqgendiff` dataset (Figure 2). It is only the fourth principle component in the `seqgendiff` dataset that seems to capture the group membership (Supplementary Figure S3). Though this unwanted variation exists, with such a large sample size voom-limma can accurately estimate the effects (Supplementary Figure S4). The voom plots (visualizing the mean-variance trend [Law et al., 2014]) are about the same in the GTEx and `seqgendiff` data, but the distribution of the square-root standard deviations appears more symmetric in the `powsimR` dataset (Figure 3). There is also an uncharacteristic hook in the mean-variance trend in the `powsimR` dataset for low-counts. These visualizations indicate that `seqgendiff` can generate more realistic datasets for RNA-seq simulation.

**Figure 1:**
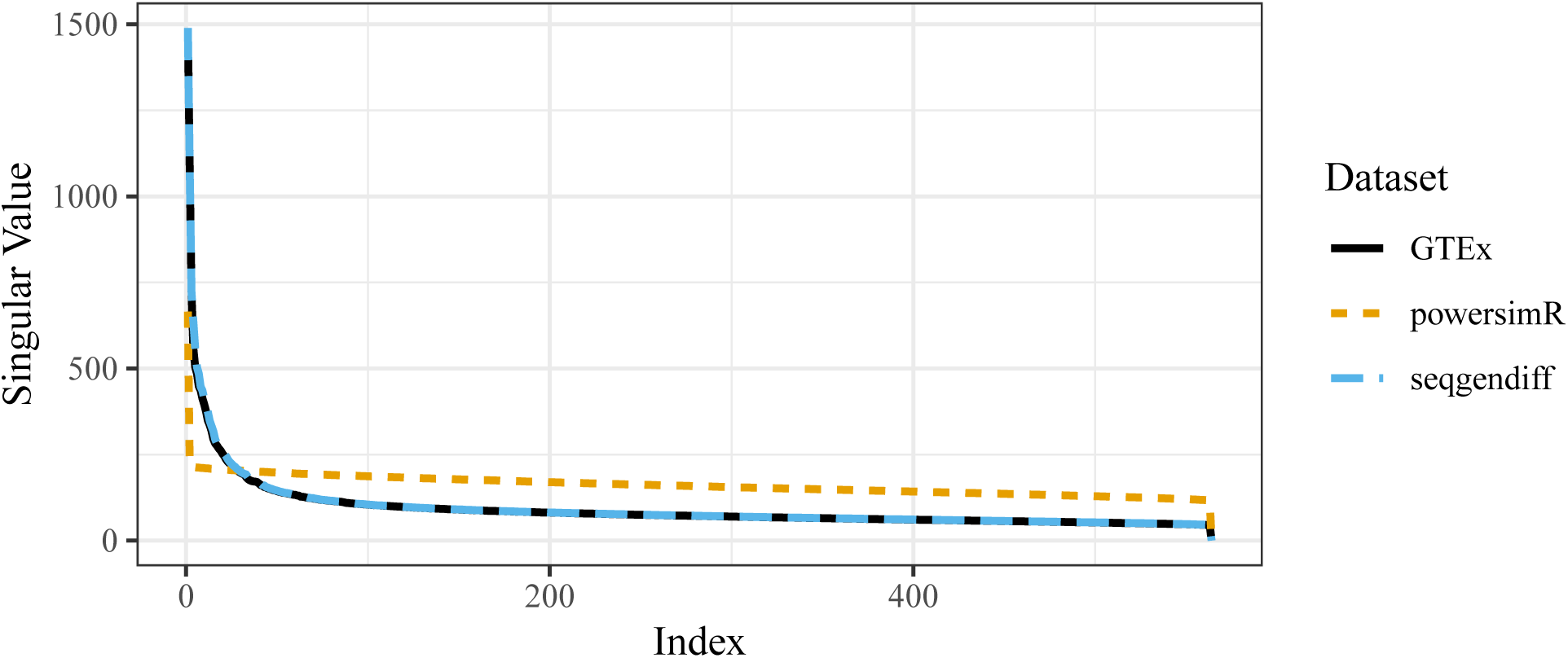
Scree plots for the GTEx dataset (black), `powsimR` dataset (orange), and the `seqgendiff`dataset (blue). The singular values for the GTEx and `seqgendiff` datasets are almost identical.

**Figure 2:**
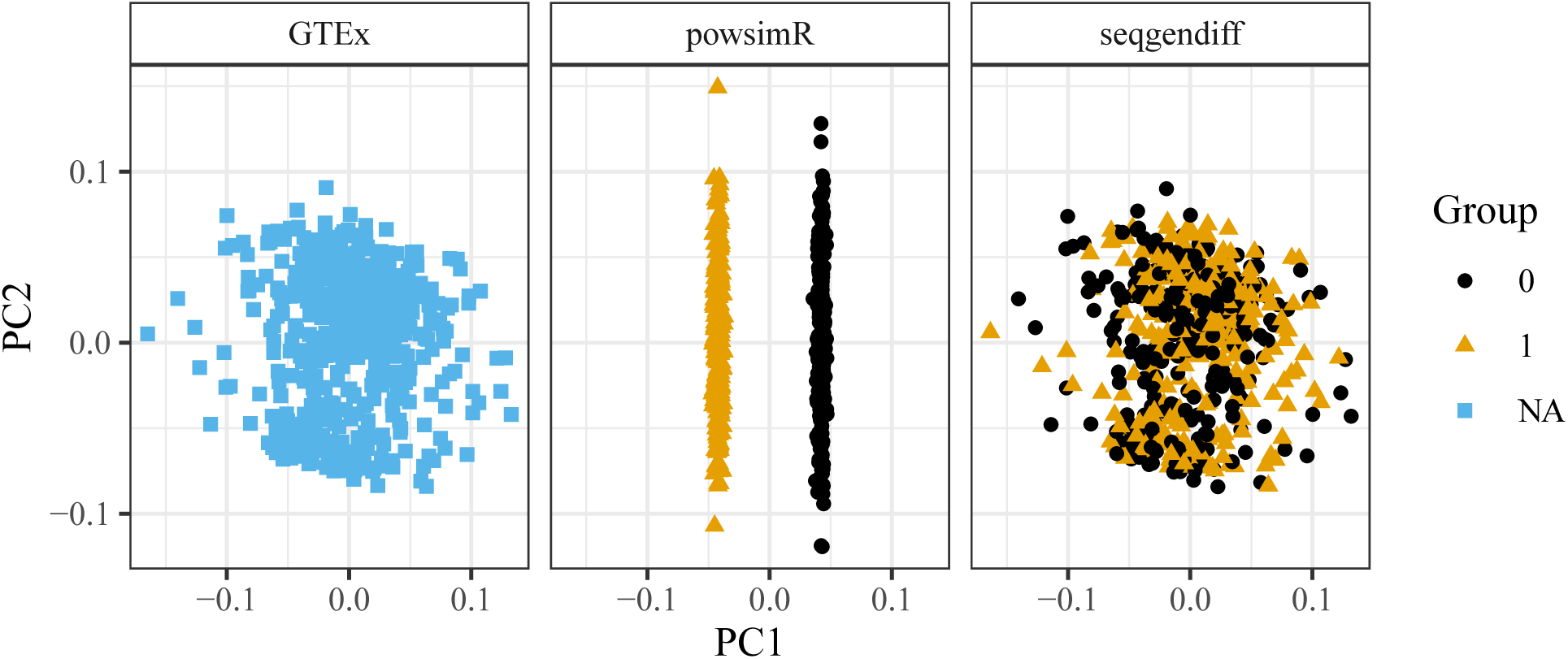
First and second principle components for the GTEx dataset (left), the `powsimR` dataset (center), and the `seqgendiff` dataset (right).

**Figure 3:**
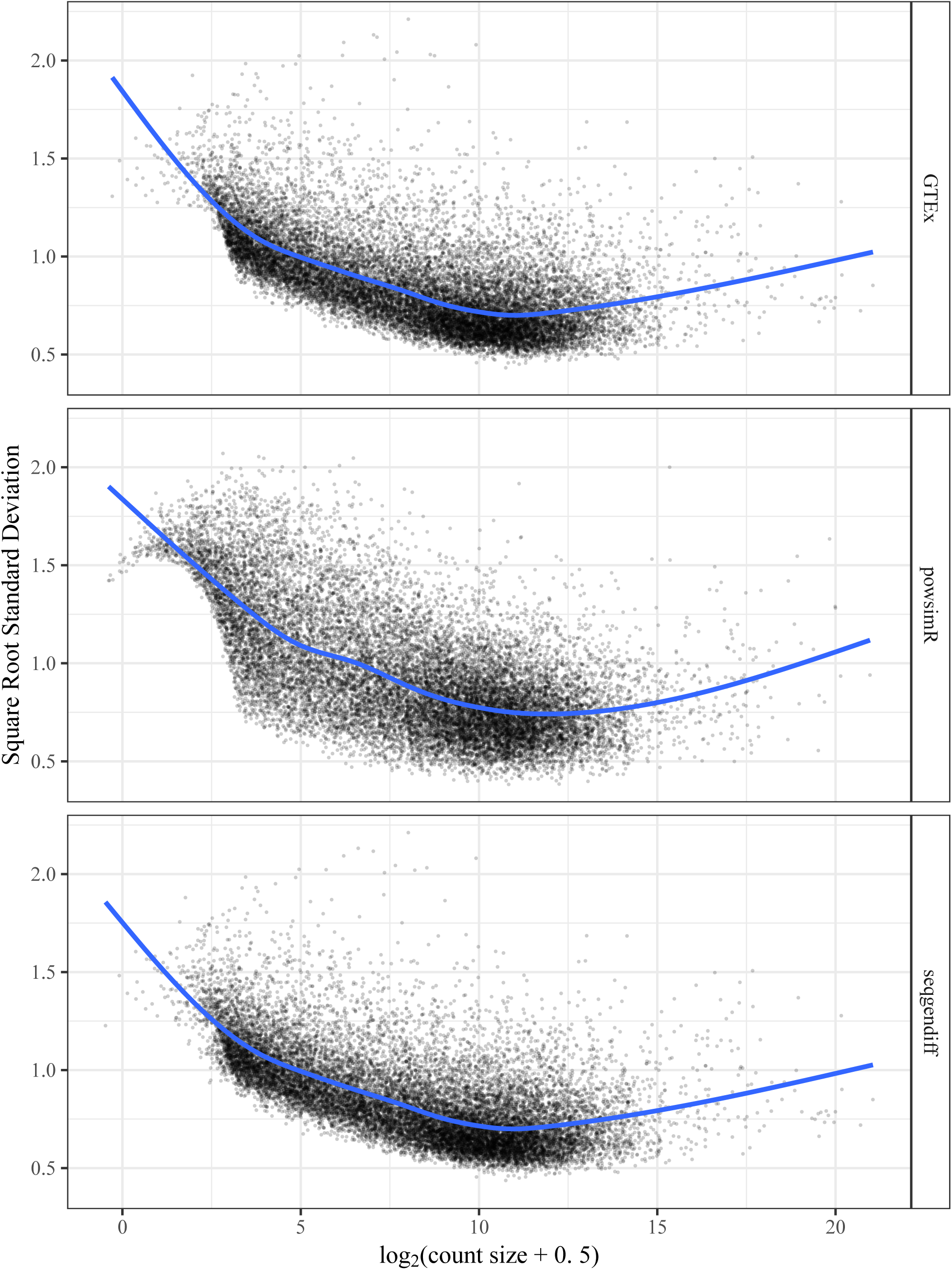
Voom plots [Law et al., 2014] visualizing the mean-variance trend in RNA-seq datasets. The voom plots are visually similar for the GTEx and `seqgendiff` datasets. The `powsimR` dataset has an uncharacteristic hook near the low counts in its voom plot.

### 3.2 Effects on Differential Expression Analysis Simulations

The differences in real versus simulated data (as discussed in Section 3.1) have real implications when evaluating methods in simulation studies. To demonstrate this, we used the GTEx muscle data to simulate RNA-seq data from the two-group model as in Section 3.1. We did this for *N* = 10 individuals, *G* = 10,000 genes, setting 90% of the genes to be null, and generating the log_2_-fold change from a *N* (0, 0.8^2^) distribution for the non-null genes. We simulated 500 datasets this way using both `seqgendiff` and `powsimR`. Each replication, we applied DESeq2 [Love et al., 2014], edgeR [Robinson et al., 2009], and voom-limma [Law et al., 2014] to the simulated datasets. We evaluated the methods based on (i) false discovery proportion when using Benjamini-Hochberg [Benjamini and Hochberg, 1995] to control false discovery rate at the 0.05 level, (ii) power to detect non-null effects based on a 0.05 false discovery rate control threshold, and (iii) mean squared error of the estimates.

We wanted to make sure that the datasets generated from `powsimR` and `seqgendiff` were comparable, so we measured the proportion of variance explained (PVE) by the group membership for each gene, which we define as

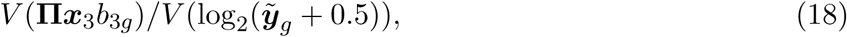

where *b*_3*g*_ is log_2_-fold change for gene 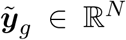 is the *g*th row of 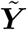, and *V* (·) returns the empirical variance of a vector. When we looked at the median (over the non-null genes) PVE across the datasets, the `seqgendiff` datasets and `powsimR` datasets had the same median PVE on average, though there was higher variability in the median PVE among the `seqgendiff` datasets (Supplementary Figure S5).

Boxplots of the false discovery proportion for each method in each dataset can be found in Figure 4. Both the `powsimR` and `seqgendiff` datasets indicate that only voom-limma can control false discovery rate adequately at the nominal level. However, the results based on the `seqgendiff` datasets indicate that there is a lot more variability in false discovery proportion than indicated by the `powsimR` datasets. In particular, it does not seem uncommon for `seqgendiff` to generate datasets with false discovery proportions well above the nominal rate. If a researcher were using only the theoretical datasets generated by `powsimR`, they would be overly confident in the methods’ abilities to control false discovery proportion. Supplementary Figure S6 also indicates that methods generally have much more variable power between the `seqgendiff` datasets than between the `powsimR` datasets. Interestingly, the `seqgendiff` datasets indicate that methods tend to have smaller mean squared error than indicated by the `powsimR` datasets (Supplementary Figure S7).

**Figure 4:**
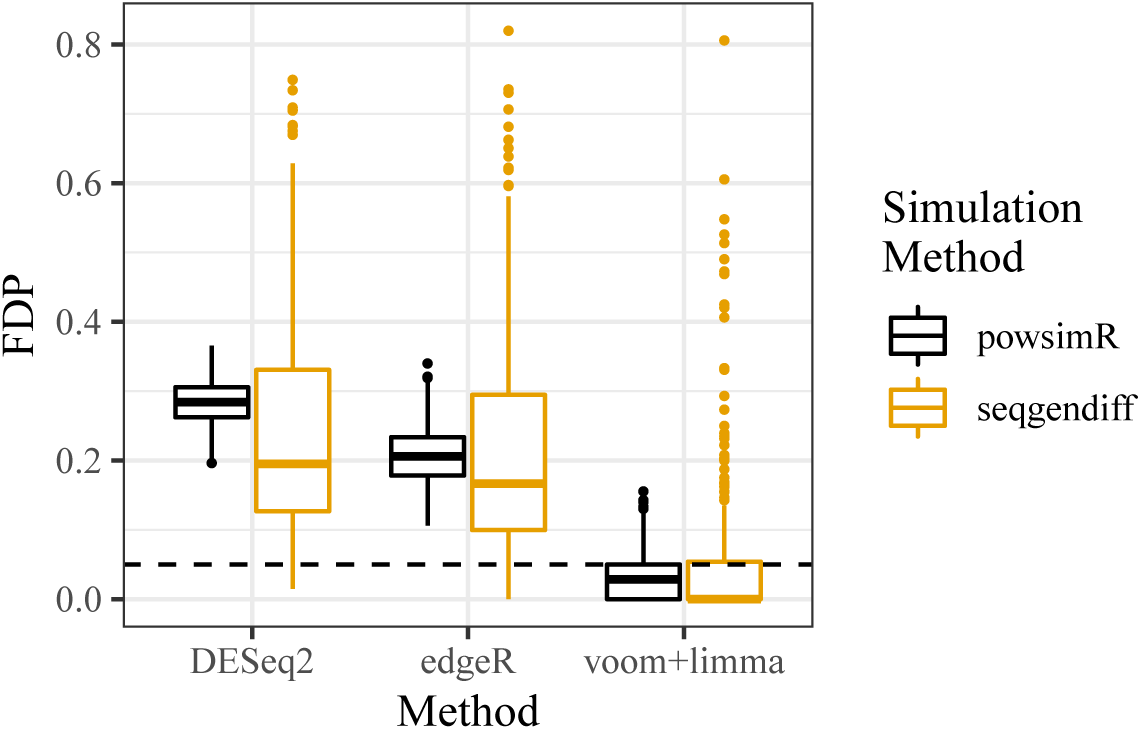
Boxplots of false discovery proportion (FDP) (*y*-axis) for various differential expression analysis methods (*x*-axis) when applied on different simulated datasets (color). Benjamini-Hochberg was used to control for false discovery rate at the 0.05 level (horizontal dashed line).

### 3.3 Evaluating Factor Analyses

As we hope we have made clear, there are many approaches to differential expression analysis (Section 2.2), confounder adjustment (Section 2.3), library size adjustment (Section 2.4), and factor analysis (Section 2.5). We believe it to be beyond the scope of this work to exhaustively evaluate all of these methods — especially since new methods are being developed each year. Rather, we hope our simulation procedures will be used by the research community to more realistically evaluate and benchmark their approaches to RNA-seq data analysis.

However, as a final highlight to the utility of our simulation approaches, we demonstrate these simulation techniques in one application: evaluating factor analysis methods in RNA-seq (Section 2.5). We have chosen to highlight this particular application because it uses the more general simulation techniques beyond the two-group model, which were first demonstrated in [Gerard and Stephens, 2017].

We chose to focus on the following methods based on (i) previous use in expression studies, (ii) software availability, (iii) popularity, and (iv) ease of use.

1. Principle component analysis (PCA) [Hotelling, 1933],
2. Sparse singular value decomposition (SSVD) [Yang et al., 2014],
3. Independent component analysis (ICA) [Hyvärinen and Oja, 2000],
4. Factors and loadings by adaptive shrinkage (*flash*), an empirical Bayes matrix factorization approach proposed in [Wang and Stephens, 2018], and
5. Probabilistic estimation of expression residuals (PEER) [Stegle et al., 2010], a Bayesian factor analysis used in the popular PEER software to adjust for hidden confounders in gene expression studies.

All factor analysis methods were applied to the log_2_-counts after adding half a pseudo-count. To simulate RNA-seq data, we took the muscle GTEx data [GTEx Consortium, 2017] and removed all genes with less than an average of 10 reads per sample. Each replicate, we added a rank-1 term. That is we assumed model (9) for the muscle GTEx data, then generated RNA-seq data such that

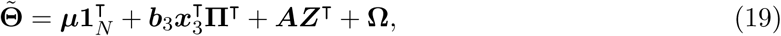

where we simulated the components of ***x***_3_ and the non-zero components of ***b***_3_ from independent normal distributions. We varied the following parameters of the simulation study:

1. The sample size: *N* ∈ {10, 20, 40}
2. The signal strength: the standard deviation of the loadings (the 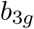’s) was set to one of {0.4, 0.8}, with higher standard deviations corresponding to higher signal. These values were chosen to have the median PVE vary greatly between the two settings (Supplementary Figure S8),
3. The sparsity: the proportion of loadings (the 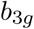’s) that are 0 was set to one of {0, 0.9}, and
4. The target correlations of the added factor with the first unobserved factor: *r* ∈ {0, 0.5}.

This resulted in 24 unique simulation parameter settings. We also used 1000 genes each replication. For each setting, we ran 100 replications of generating data from model (19), and fitting the factors with the five methods under study assuming model (11) after we estimated the number of hidden factors using parallel analysis [Buja and Eyuboglu, 1992].

We chose three metrics to evaluate the performance of the different factor analysis methods:

1. The minimum mean squared error between **Π*x***_3_ and the columns of 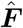. To account for scale and sign unidentifiability, the estimated factors and the added factor were all scaled to have an *ℓ*^2^-norm of 1 prior to calculating the mean squared error. This measure is meant to evaluate if any of the estimated factors corresponds to the added factor.
2. The minimum mean squared error between ***b***_3_ and the columns of 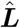. We again accounted for scale and sign unidentifiability by calculating the mean squared error after scaling the estimated and true loadings to have an *ℓ*^2^-norm of 1.
3. The angle between **Π*x***_3_ and its projection onto the column space of 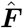. This measure is meant to evaluate if the estimated factor matrix includes **Π*x***_3_ among its unidentified factors.

The results are presented in Supplementary Figures S9-S14. Based on these figures, we have the following conclusions:

1. PEER performs very poorly when either the sparsity is high or when there are few samples. It also performs less well when the factors are correlated. A possible explanation is that PEER assumes a normal distribution on the factors and loadings, which is violated in the high-sparsity regime and is observed in the low-sparsity regime. Though, this does not explain its poor performance in small sample size settings.
2. SSVD estimates the loadings very poorly in low-sparsity regimes. This is to be as expected as SSVD assumes sparsity on the loadings. Surprisingly, though, it outperforms PCA in high sparsity regimes only when both the sample size and signal are also large.
3. ICA performs very poorly in low sparsity regimes. This is to be as expected as the normal distributions placed on the factors and loadings are a worst-case scenario for ICA. However, there is no scenario where ICA performs significantly better than PCA.
4. *flash* performs adequately in all scenarios and performs best in high-sparsity and high-signal regimes.
5. PCA performs adequately in most scenarios, and is only truly outperformed in high sparsity high signal regimes.
6. Based on these initial explorations, we would recommend users not use PEER, SSVD, or ICA and instead try either PCA or *flash*.

## 4 Discussion

We have focused on a log-linear model because of the large number of applications this generates (Sections 2.2, 2.3, 2.4, and 2.5). This linearity (on the log-link scale) is represented by the structure of the ***Q*** matrix in Procedure 1. However, it is possible to replace ***Q*** by any arbitrary *G* × *N* matrix that has non-positive entries. This might be useful for simulations that study adjusting for non-linear effects, such as bias due to GC content [Risso et al., 2011], as it allows you to introduce non-linear effects into an RNA-seq dataset. However, these non-linear effects would still be present only on the log-scale.

Our simulation procedures may be applicable beyond evaluating competing methods. Vieth et al. [2017] used their simulation software to estimate power given the sample size in a differential expression analysis, and thus to develop sample size suggestions. Our simulation methods may be used similarly. Given a large RNA-seq dataset (such as the GTEx data used in this paper), one can repeatedly down-sample the number of individuals in the dataset and explore how sample size affects the power of a differential expression analysis.

Similarly, Robinson and Storey [2014] already demonstrated that binomial thinning may be used for sequencing depth suggestions. That is, a researcher may repeatedly thin the libraries of the samples in an large RNA-seq dataset and explore the effects on power, thereby providing sequencing depth suggestions. Unlike Robinson and Storey [2014], which does this subsampling uniformly over all counts, we allow researchers to explore the effects of heterogeneous subsampling (as in Section 2.4). This might be useful if, say, researchers have more individuals in one group than in another and so wish to explore if they can sequence the larger group to a lower depth without affecting power.

In this manuscript, we have discussed our simulation techniques in the context of RNA-seq. However, our techniques would also be applicable to the comparative analysis of metagenomics methods [Jonsson et al., 2016]. Instead of quantifying gene expression, metagenomics quantifies gene abundances within metagenomes. Our simulation techniques could be applied in this context by taking a real metagenomics dataset and adding signal to it by binomial thinning.

## 5 Conclusions

We developed a procedure to add a known amount of signal to any real RNA-seq dataset. We only assume that this signal comes in the form of a generalized linear model with a log-link function from a very flexible distribution. We demonstrated how real data contain features that are not captured by simulated data, and that this can cause important differences in the results of a simulation study. We highlighted our simulation approach by comparing a few popular factor analysis methods. We found that PCA and *flash* had the most robust performances across a wide range of simulation settings.

## Availability of data and materials

The simulation methods discussed in this paper are implemented in the `seqgendiff` R package, available on the Comprehensive R Archive Network: https://cran.r-project.org/package=seqgendiff. All code to reproduce the simulation and analysis results is available on GitHub: https://github.com/dcgerard/reproduce_fasims.

The datasets analyzed during the current study are available in the GTEx portal: https://gtexportal.org.

## Acknowledgments

We would like to thank Matthew Stephens for providing comments on a draft of this manuscript, and Joyce Hsiao for testing an early version of the `seqgendiff` software.

All graphics were made using ggplot2 [Wickham, 2016] in the R statistical language [R Core Team, 2019].

## S1 S1 Theoretical Considerations

### S1. Target Correlation

#### Theorem 1.

*Let* (*u*_*i*_, *z*_*i*_) *be iid jointly standard normal with correlation ρ for i* = 1, 2, …, *n*. *Let w*_*j*_ *be iid standard normal for j* = 1, 2, …, *n*. *Suppose we match the w*_*j*_*’s onto the u*_*i*_*’s by order statistics, resulting in* (*w*_*i*_, *u*_*i*_) *pairs such that the rank of w*_*i*_ *is the same as the rank of u*_*i*_.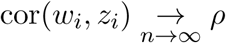.

#### Proof.

For a fixed proportion *p*, we note that *u*_(*⌈np⌉*)_ and *w*_(*⌈np⌉*)_ converge in probability to the theoretical *p*-quantile of the standard normal distribution [Arnold et al., 1992, e.g.]. Since the order statistics converge to the same values, and we match by order statistics, this implies that 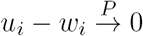. Thus, by Slutsky’s theorem, we have that 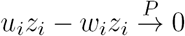.

We note that |*u*_*i*_*z*_*i*_ *- w*_*i*_*z*_*i*_| *≤* |*u*_*i*_*z*_*i*_| + |*w*_*i*_*z*_*i*_|, by the triangle inequality. The term on the right has finite expectation as (using Cauchy-Schwarz)

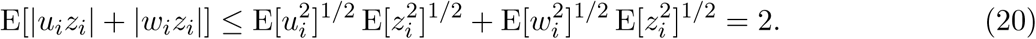

Thus, by the Lebesgue dominated convergence theorem, we have E[|*u*_*i*_*z*_*i*_ *− w*_*i*_*z*_*i*_|] → 0. Since − |*u*_*i*_*z*_*i*_ *− w*_*i*_*z*_*i*_| *≤ u*_*i*_*z*_*i*_ *− w*_*i*_*z*_*i*_ *≤* |*u*_*i*_*z*_*i*_ *− w*_*i*_*z*_*i*_|, this implies that E[*u*_*i*_*z*_*i*_] *− E*[*w*_*i*_*z*_*i*_] → 0, and the theorem is proved.□

To place the results of Theorem 1 in context of the matching in Procedure 2, note that the *u*_*i*_’s are the elements of ***U***, the *z*_*i*_’s are the elements of 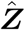, the *w*_*i*_’s are the elements of ***X***_3_, *ρ* is the target correlation between the one column in ***X***_3_ and the one column in 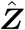, and **Π** is the permutation matrix that results in the matching of the *w*_*i*_’s and the *u*_*i*_’s.

The results of Theorem 1 can be generalized to non-standard normal distributions by appealing to the weak law of large numbers and Slutsky’s theorem.

### S1.2 Generalizing the Poisson Assumption

For simplicity, we stated a Poisson distribution as the modeling assumption in (1). However, our methods are equally valid under more general conditions. We begin by showing how our methods are valid when using the negative binomial distribution, which is perhaps the most common distribution used to analyze RNA-seq counts [Robinson and Smyth, 2007a,b, Love et al., 2014]. To see this, we prove the following simple lemma which, though less well-known than Lemma 1, can still be found in some elementary texts (or at least a version of the following lemma) [exercise 4.32 of Casella and Berger, 2002, e.g.].

#### Lemma 2.

*Suppose y* ∼ NB(*µ, ϕ*), *where we are using the parameterization such that* E[*y*] = *µ and* var(*y*) = *µ*(1 + *µϕ*). *Also suppose that* 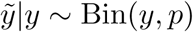. *Then* 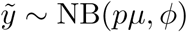.

#### Proof.

Using the hierarchical characterization of the negative binomial distribution, we have that

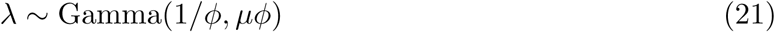

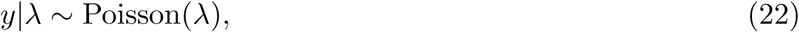

where 1*/ϕ* is the shape parameter and *µϕ* is the scale parameter. This implies that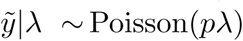. But *pλ* ∼ Gamma(1*/ϕ, pµϕ*) by elementary properties of the gamma distribution. Hence, by the hierarchical characterization of the negative binomial distribution, we have that 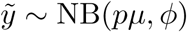.□

The zero-inflated negative binomial distribution is sometimes used to model single-cell RNA-seq data as it can account for the abundance of zeros observed in such data [Miao et al., 2018, Risso et al., 2018, Eraslan et al., 2019]. A random variable *y* is distributed zero-inflated negative binomial, denoted *y* ∼ ZINB(*π, µ, ϕ*), if it is generated by the following hierarchical process:

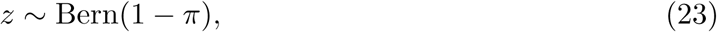

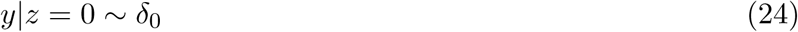

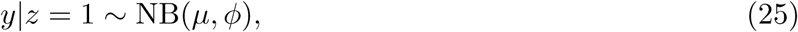

where *δ*_0_ is the degenerate distribution with a point-mass at 0. In words, the counts are either 0 with probability *π* or follow a negative binomial distribution with probability 1 *- π*. Our methods are equally valid in the zero-inflated negative binomial case.

#### Lemma 3.

*Suppose y* ∼ ZINB(*π, µ, ϕ*) *and* 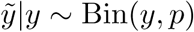. *Then* 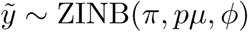.

*Proof*. It’s sufficient to note that

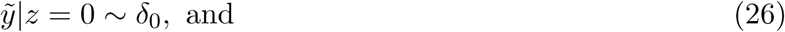

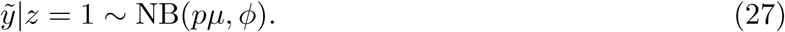

□

Finally, our simulation methods preserve the count distribution in the rich class of distributions which are mixtures of binomial and negative binomial distributions (some examples within this class of distributions are plotted in Supplementary Figure S1).

#### Lemma 4.

*Let π*_0_, *π*_1_, …*π*_*M*_ *and τ*_1_, *τ*_2_, …, *τ*_*L*_ *be non-negative mixing proportions such that*

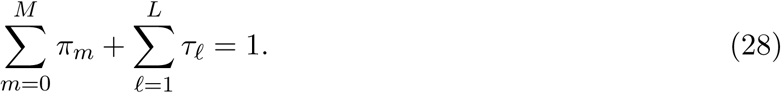

*Suppose that y has a PMF which is a mixture of binomial and negative binomial PMF’s*

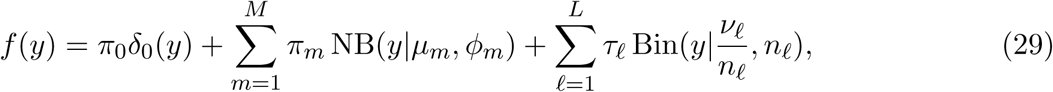

*where* NB(*y*|*µ*_*m*_, *ϕ*_*m*_) *is the negative binomial PMF with mean µ*_*m*_ *and dispersion ϕ*_*m*_, *and* Bin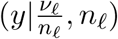 *is the binomial PMF with mean ν*_*l*_ *and success probability ν_l_/n_l_*. *Suppose that* 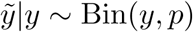. *Then*

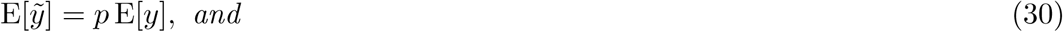

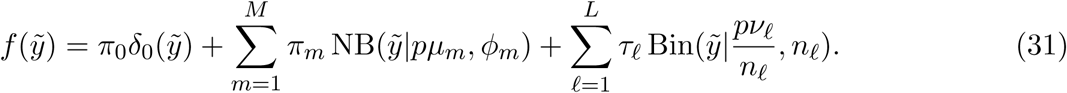

#### Proof.

Equation (30) is just a consequence of the law of total expectation. To prove (31), note that if *y* ∼ NB(*µ*_*m*_, *ϕ*_*m*_) then 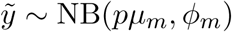 and if *y* ∼ Bin(*ν /n*, *n*) then 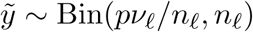. The proof follows by conditioning on the latent mixing group.

## S2 Additional Simulations

### S2.1 Correlation Estimator

We explored the effects of changing the target correlation on the true correlation. We varied the sample size, *N* ∈ {6, 10, 20}, and the target correlations between ***z*** and the two columns in **Π*X***_3_, ***r*** ∈ {(0, 0), (0.5, 0), (0.9, 0), (0.5, 0.5)}. Under each unique combination of simulation parameter settings, we iterative drew ***z*** ∈ ℝ^*N*^ from a standard normal. We also drew ***X***_3_ ∈ ℝ^*N*×2^ according to two schemes:

1. Normal: Each element of ***X***_3_ is independently drawn from a standard normal distribution, and
2. Indicator: The first column of ***X***_3_ consists of (1, 0, 1, 0, …, 1, 0)^T^, and the second column of ***X***_3_ consists of 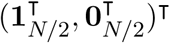.

Each replicate, we used Procedure 4 to estimate the correlation between **Π*X***_3_ and ***z***. We did this for a total of 100 replications for each combination of simulation parameters.

The results are presented in Supplementary Figure S15. Because we are approximating the expected (conditional on ***z***) Pearson correlation between the columns of **Π*X***_3_ and ***z***, the true correlations between **Π*X***_3_ and ***z*** are approximately the mean of the estimates over the 100 replications (see (17)). From Supplementary Figure S15, we note that the true correlation is generally closer to 0 than the target correlation. When the sample size is 20, there seems to be very little variability in the correlation estimates.

## S3 Supplementary Figures

**Figure S1:**
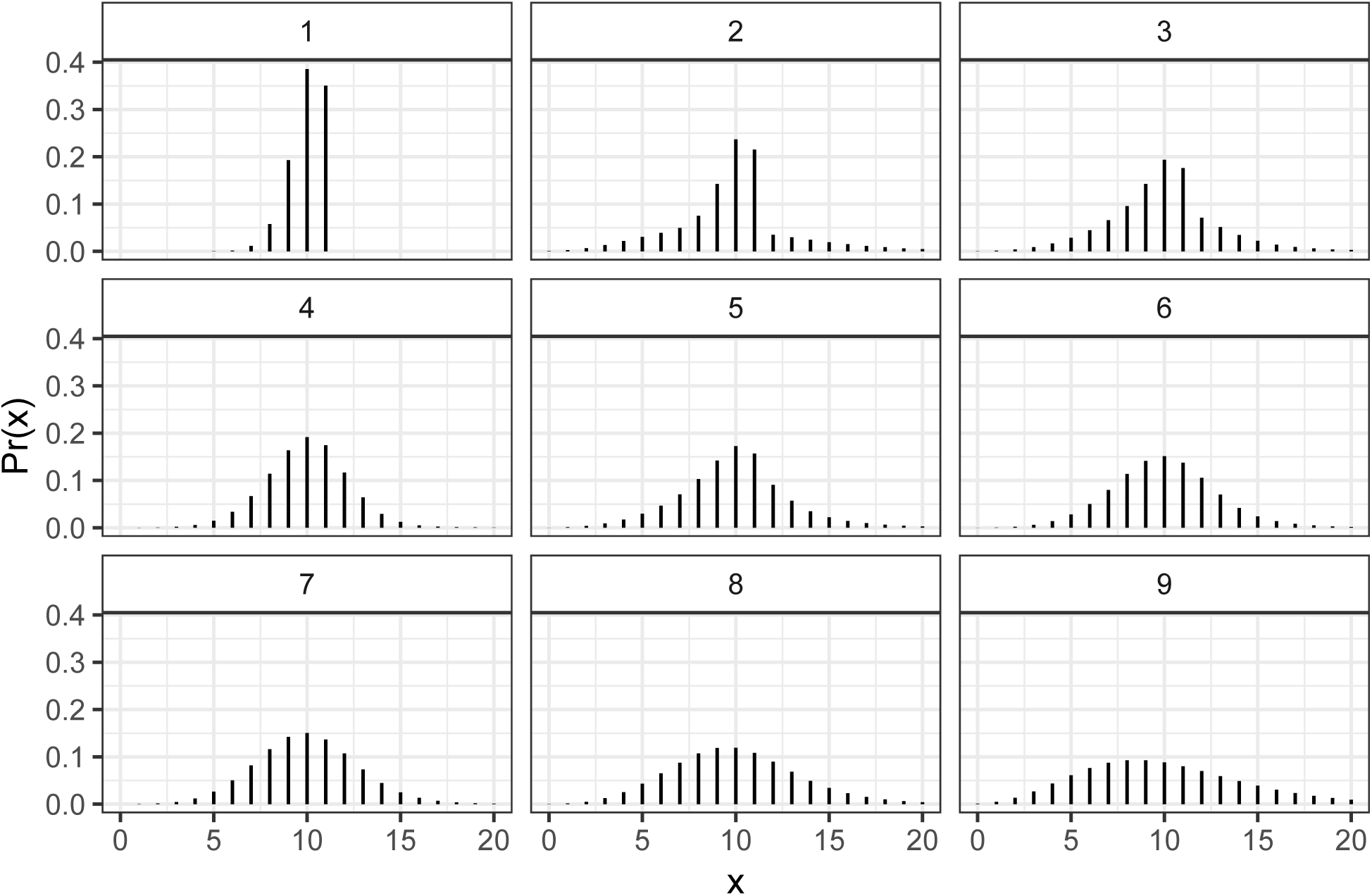
Some distributions in the class of mixtures of binomial and negative binomial distributions, demonstrating the flexibility of this class. The mean was set to 10 for all mixing components.

**Figure S2:**
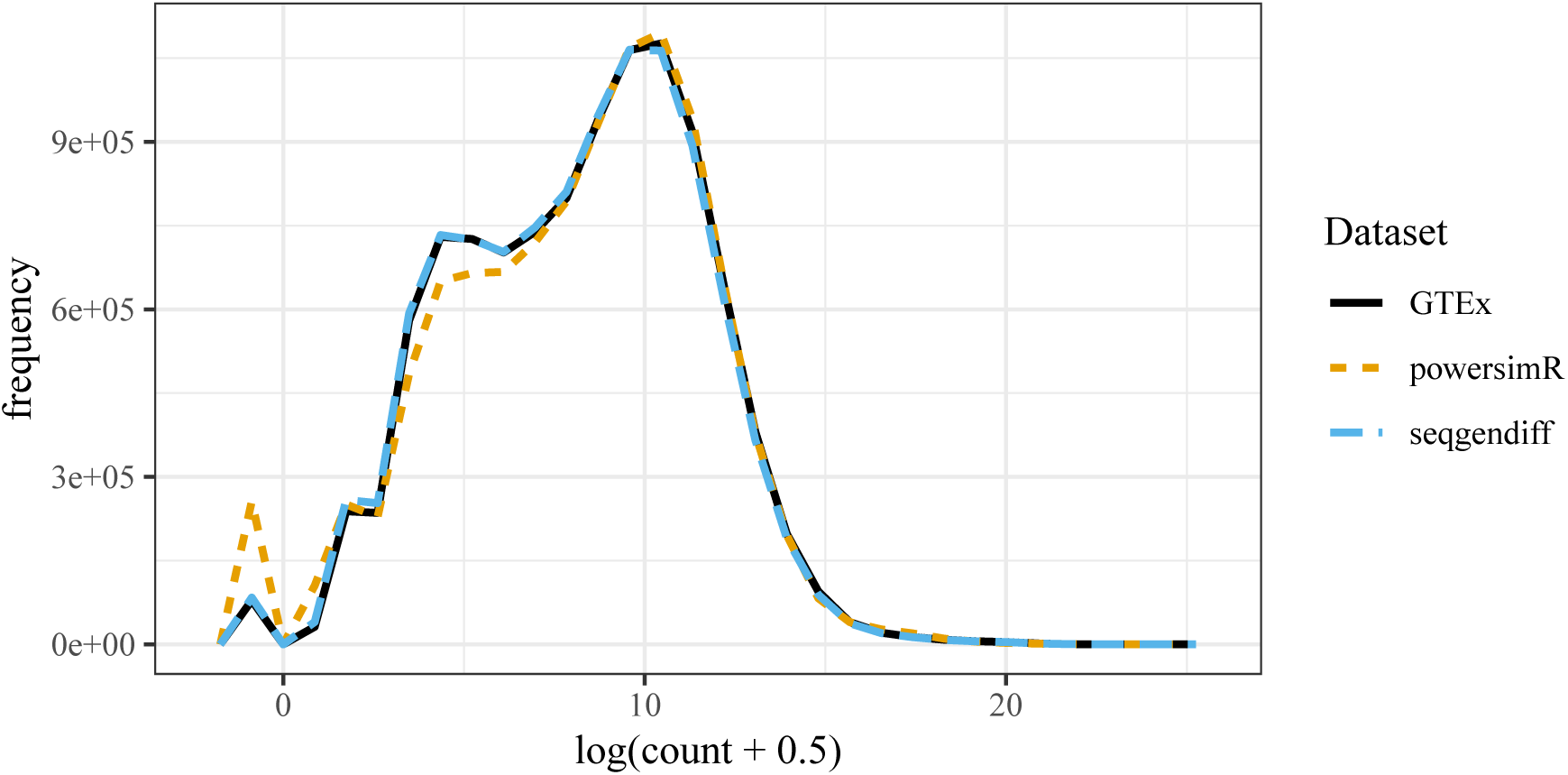
Frequency polygons for the log_2_ counts for the dataset simulated by `seqgendiff` (blue), the dataset simulated by `powsimR` (orange), and the GTEx muscle dataset (black). The `seqgendiff` and GTEx count distributions are almost exactly the same.

**Figure S3:**
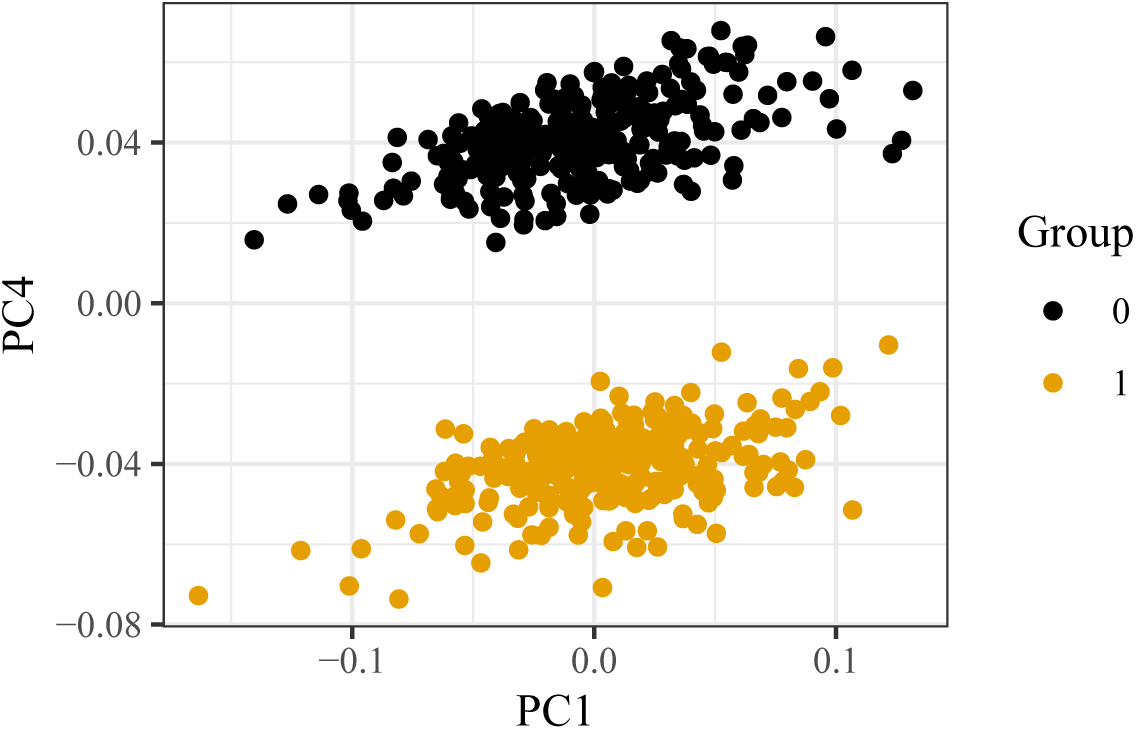
First principle component (*x*-axis) versus the fourth principle component (*y*-axis) in the `seqgendiff` dataset. The fourth principle component seems capture group membership.

**Figure S4:**
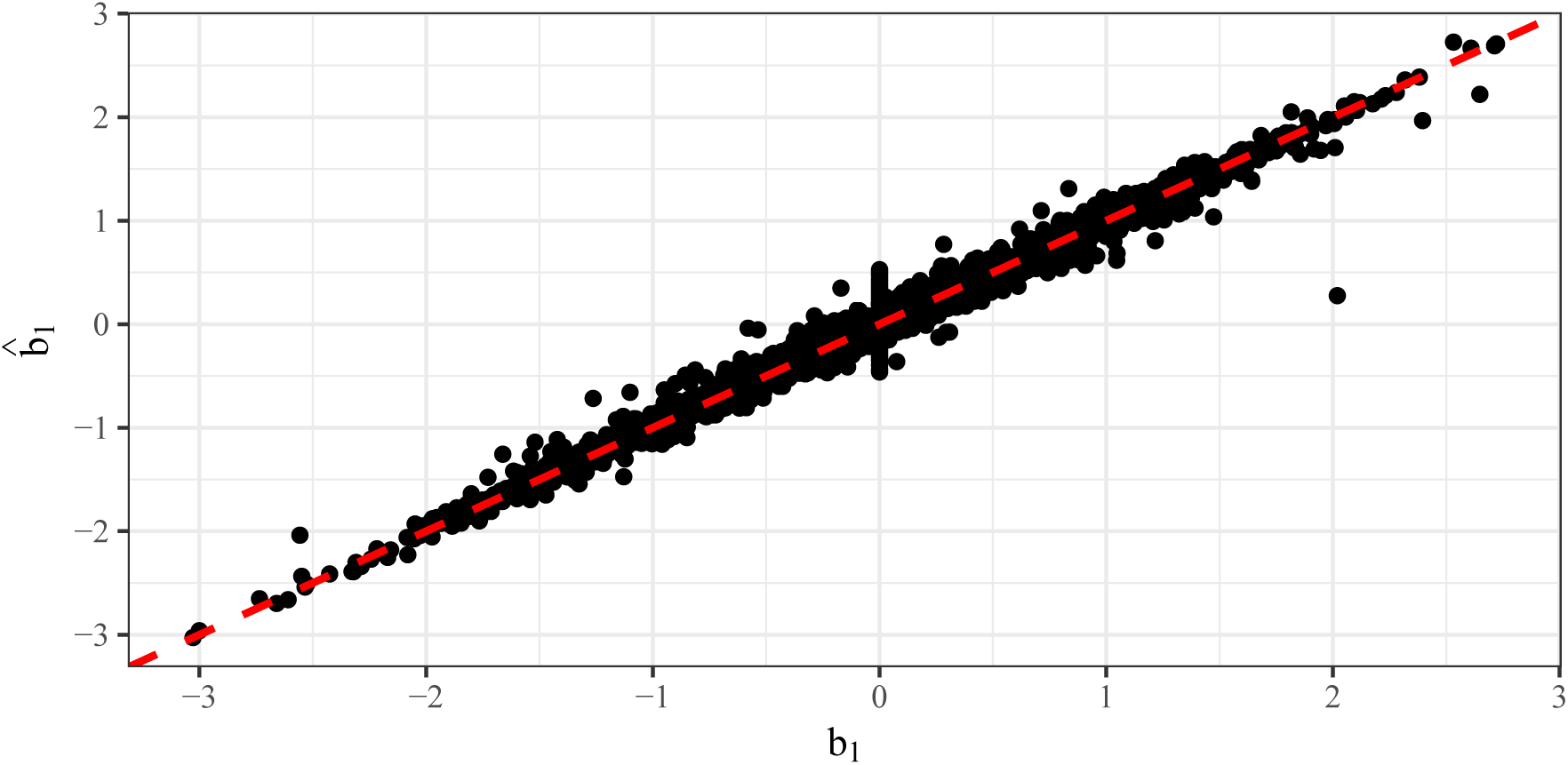
True coefficient values (*x*-axis) versus their corresponding estimates (*y*-axis) in the `seqgendiff` dataset. Estimates were obtained using the voom-limma pipeline [Smyth, 2004, Law et al., 2014].

**Figure S5:**
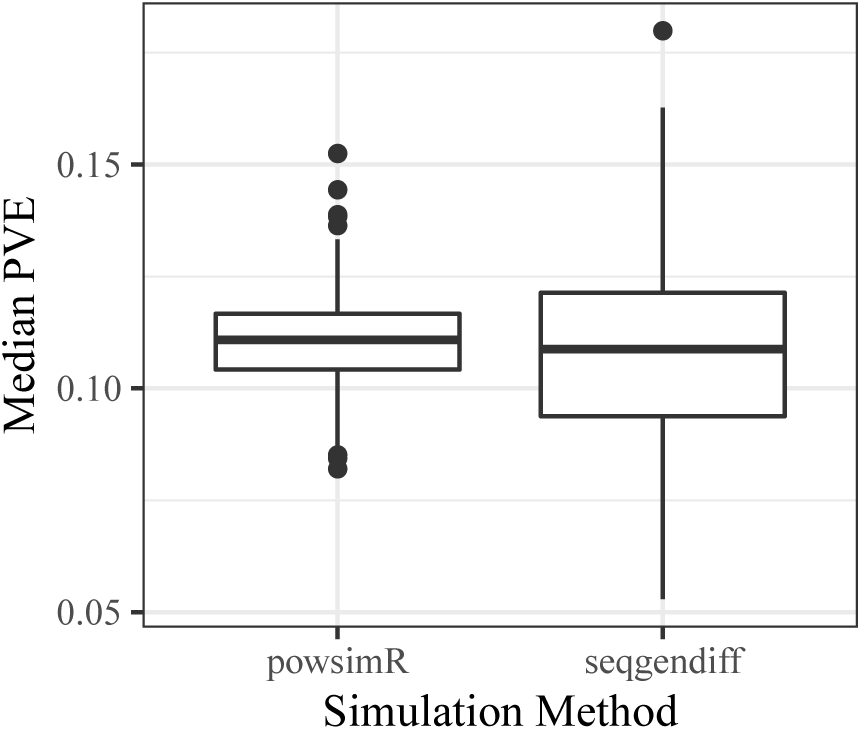
Median (across non-null genes) proportion of variance explained (18) (*y*-axis) for datasets generated by `powsimR` or `seqgendiff` (*x*-axis). The two sets of simulated datasets have the same expected median PVE, though the median PVE is more variable among the `seqgendiff` datasets.

**Figure S6:**
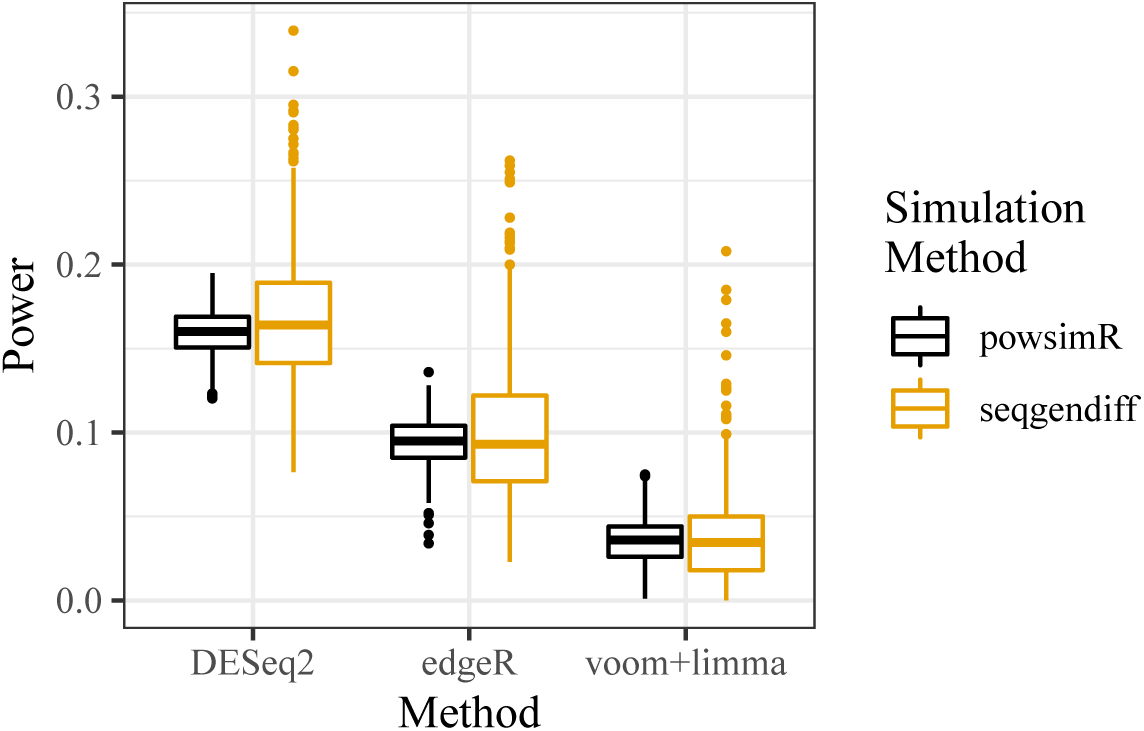
Boxplots of power (*y*-axis) for various differential expression analysis methods (*x*-axis) when applied on different simulated datasets (color).

**Figure S7:**
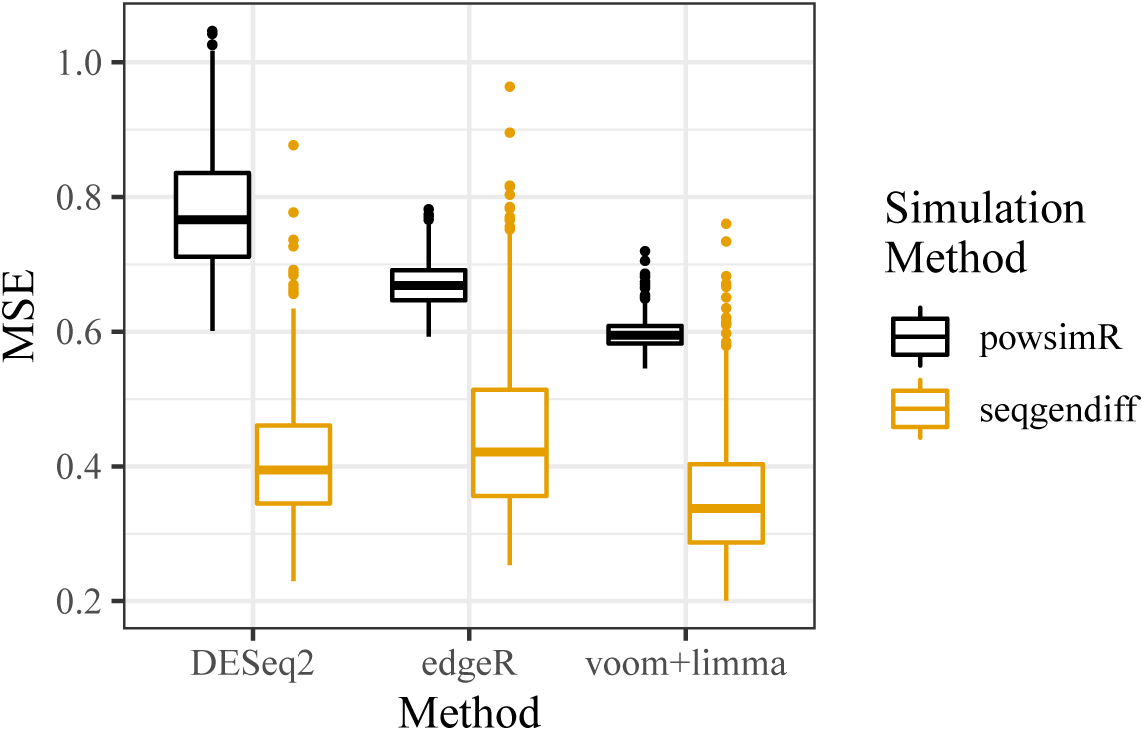
Boxplots of mean squared error (*y*-axis) for various differential expression analysis methods (*x*-axis) when applied on different simulated datasets (color).

**Figure S8:**
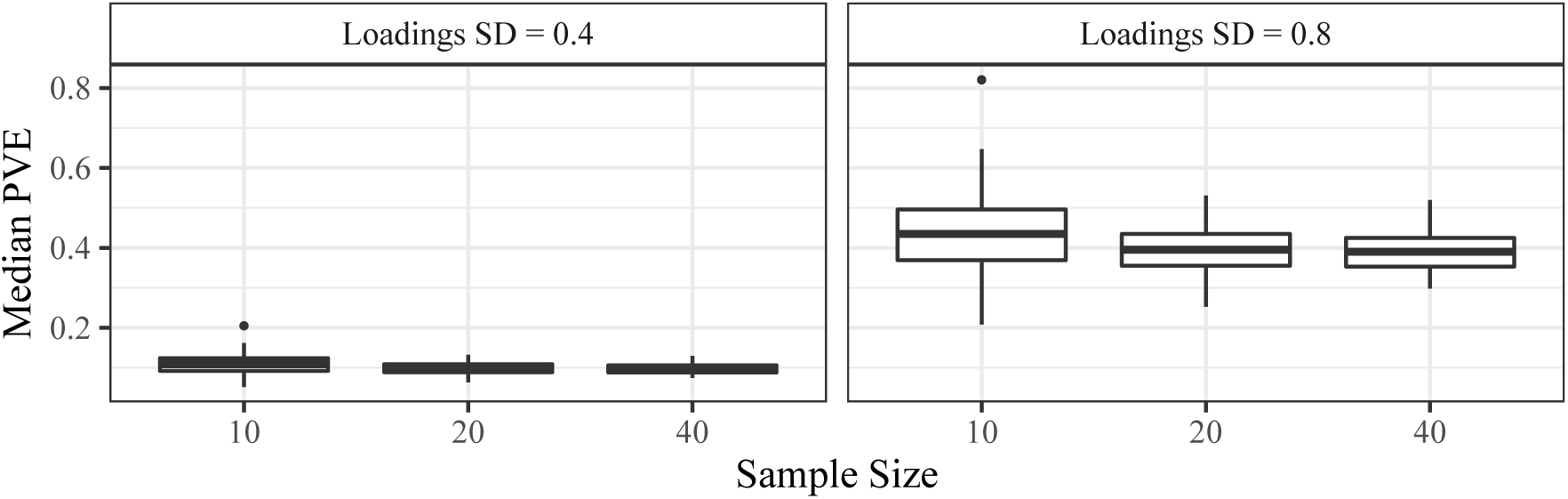
Median proportion of variance explained (18) for the genes with non-zero loadings (*y*-axis) stratified by the sample size (*x*-axis) from the simulation study in Section 3.3. The facets index the different standard deviations of the loadings.

**Figure S9:**
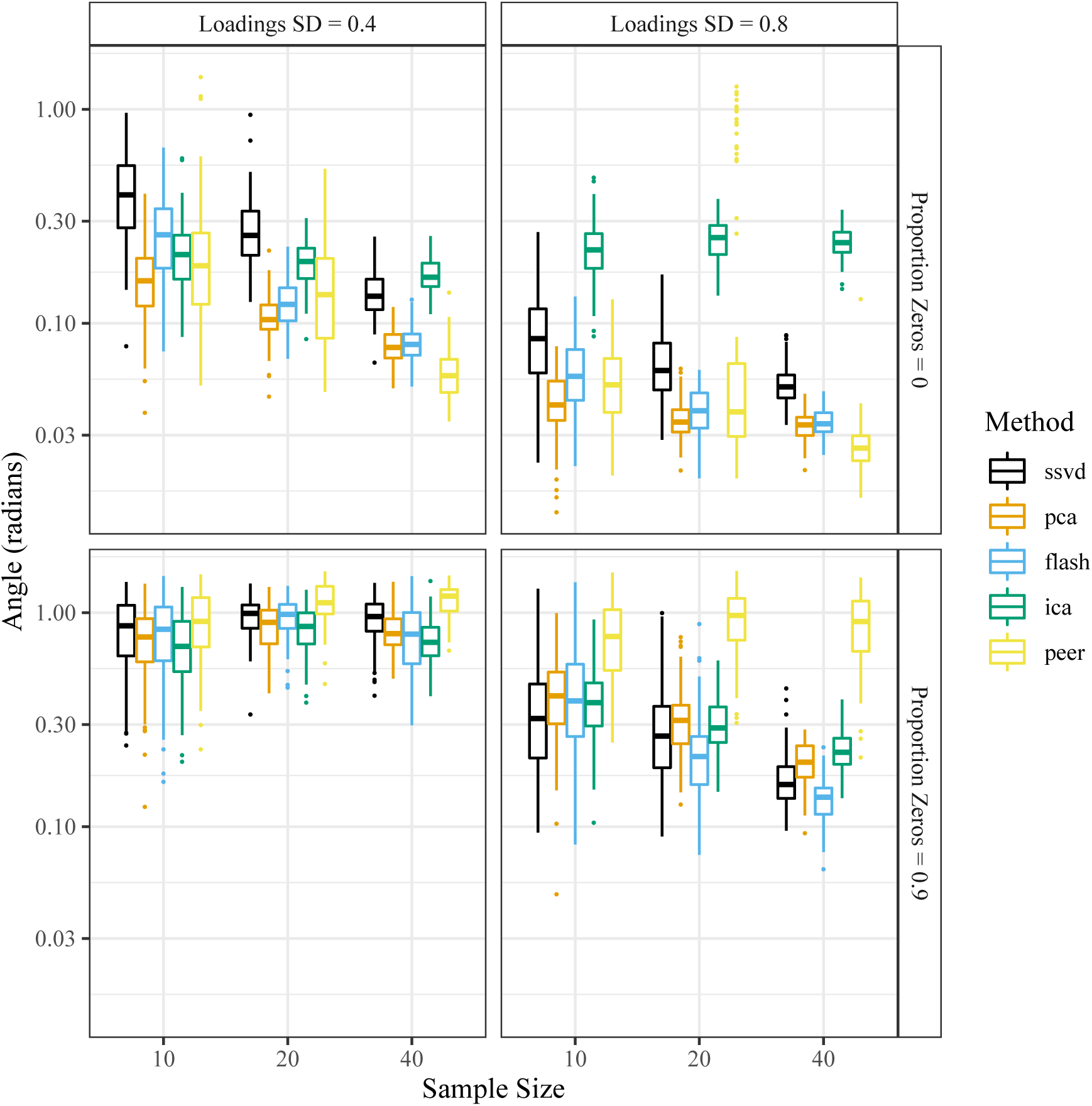
Angle (log10-scale) between the added factor and its projection onto the column space of the estimated factors (*y*-axis), stratified by sample size (*x*-axis), factor analysis method (color), signal strength (column facets), and sparsity of the loadings (row facets). The target correlation between the added factor and the first unobserved factor was set to 0. A smaller angle is better.

**Figure S10:**
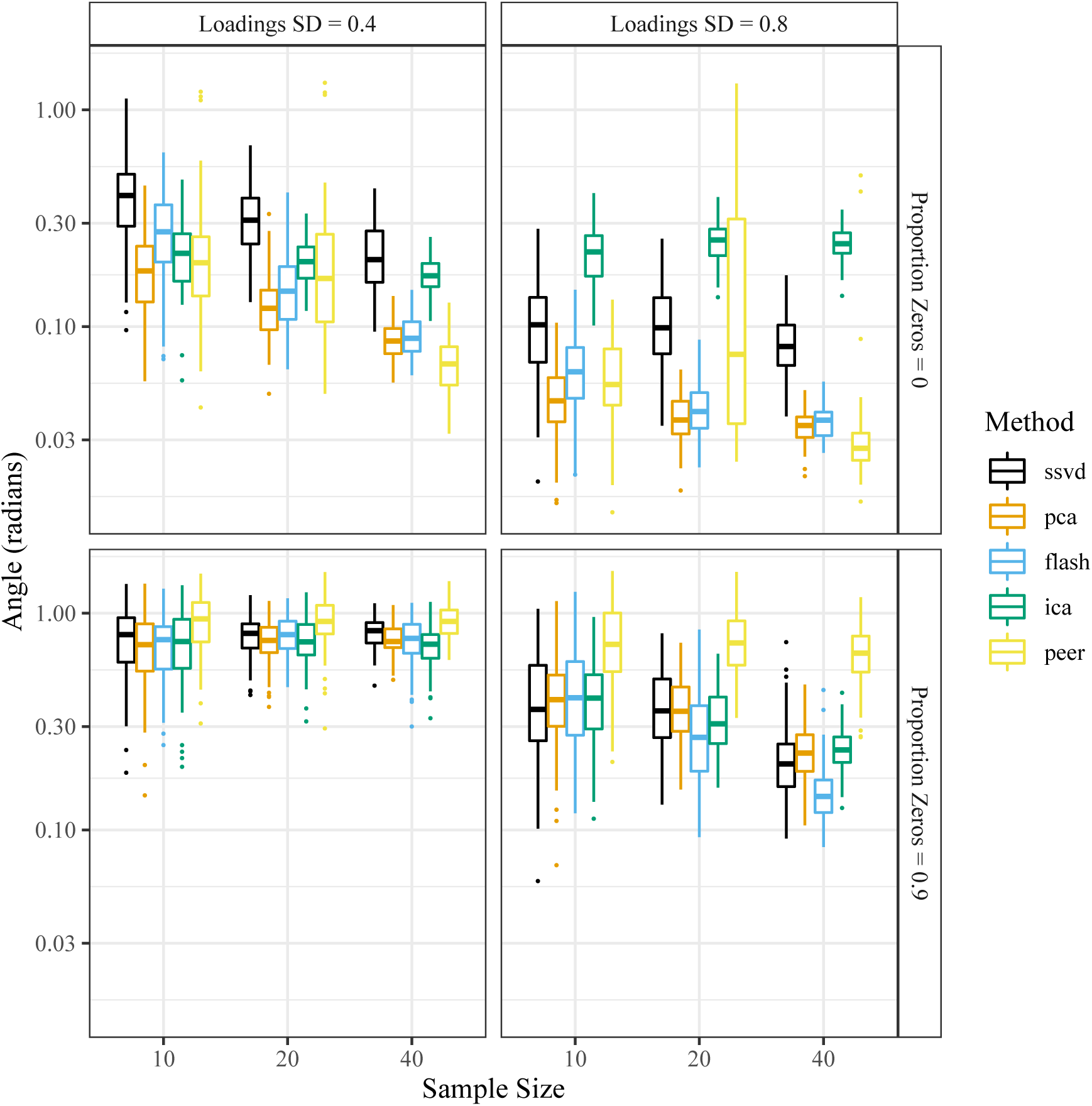
Angle (log10-scale) between the added factor and its projection onto the column space of the estimated factors (*y*-axis), stratified by sample size (*x*-axis), factor analysis method (color), signal strength (column facets), and sparsity of the loadings (row facets). The target correlation between the added factor and the first unobserved factor was set to 0.5. A smaller angle is better.

**Figure S11:**
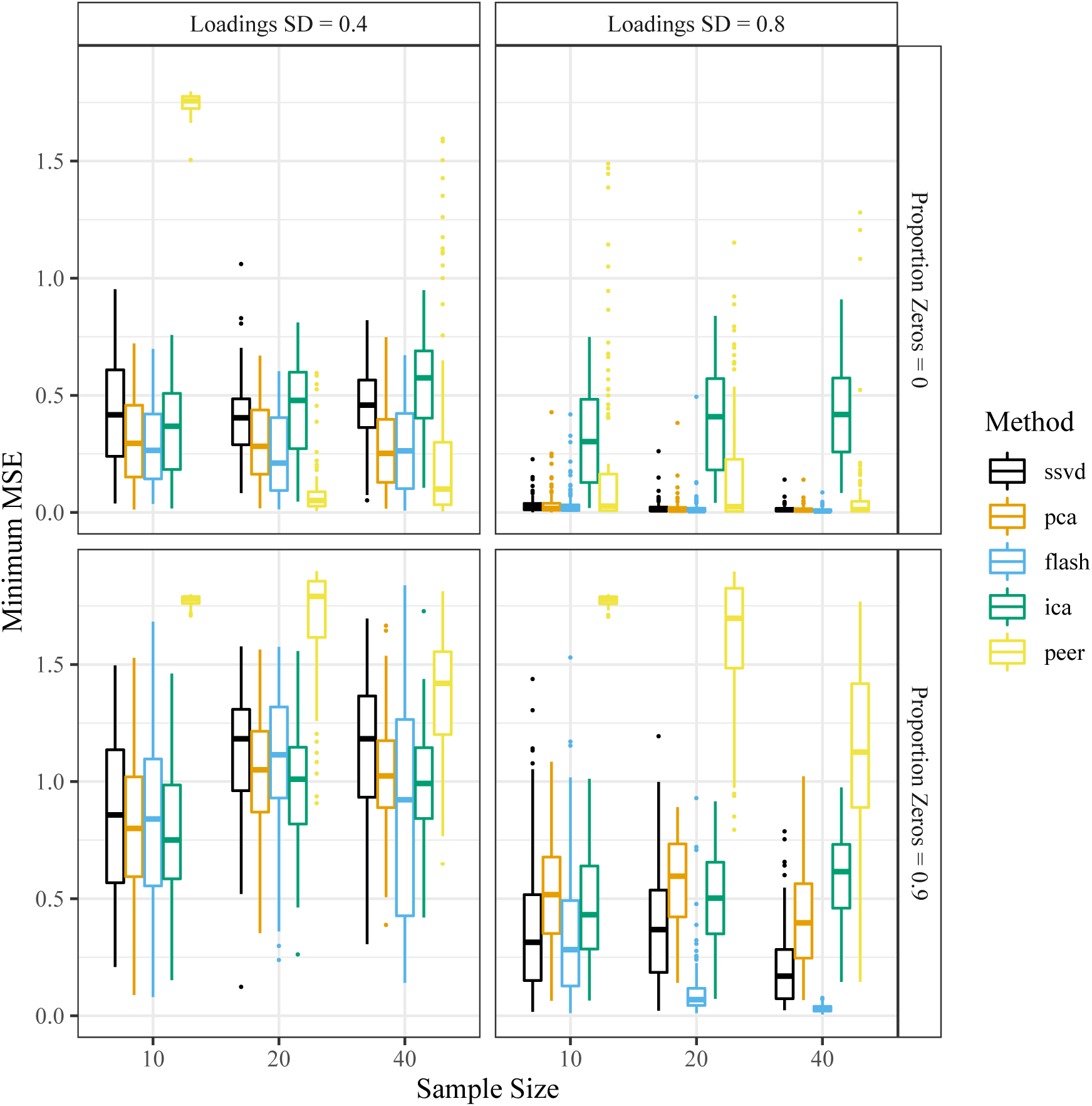
Minimum mean squared error between the added factor and the estimated factors (*y*-axis), stratified by sample size (*x*-axis), factor analysis method (color), signal strength (column facets), and of the loadings sparsity (row facets). Before calculating the MSE, all factors were scaled to have *ℓ*^2^-norm of 1. The target correlation between the added factor and the first unobserved factor was set to 0. A smaller minimum MSE is better.

**Figure S12:**
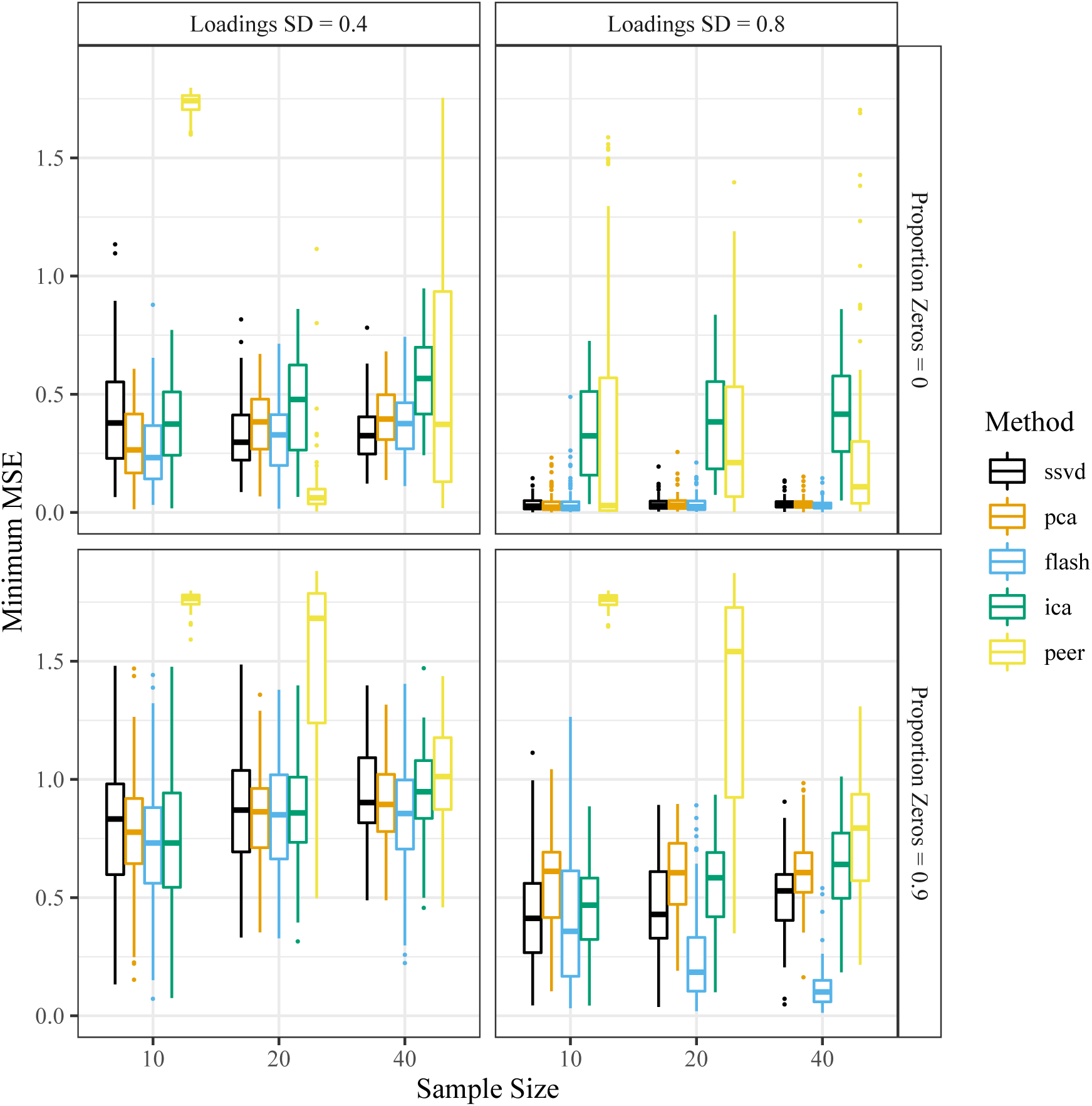
Minimum mean squared error between the added factor and the estimated factors (*y*-axis), stratified by sample size (*x*-axis), factor analysis method (color), signal strength (column facets), and sparsity of the loadings (row facets). Before calculating the MSE, all factors were scaled to have *ℓ*^2^-norm of 1. The target correlation between the added factor and the first unobserved factor was set to 0.5. A smaller minimum MSE is better.

**Figure S13:**
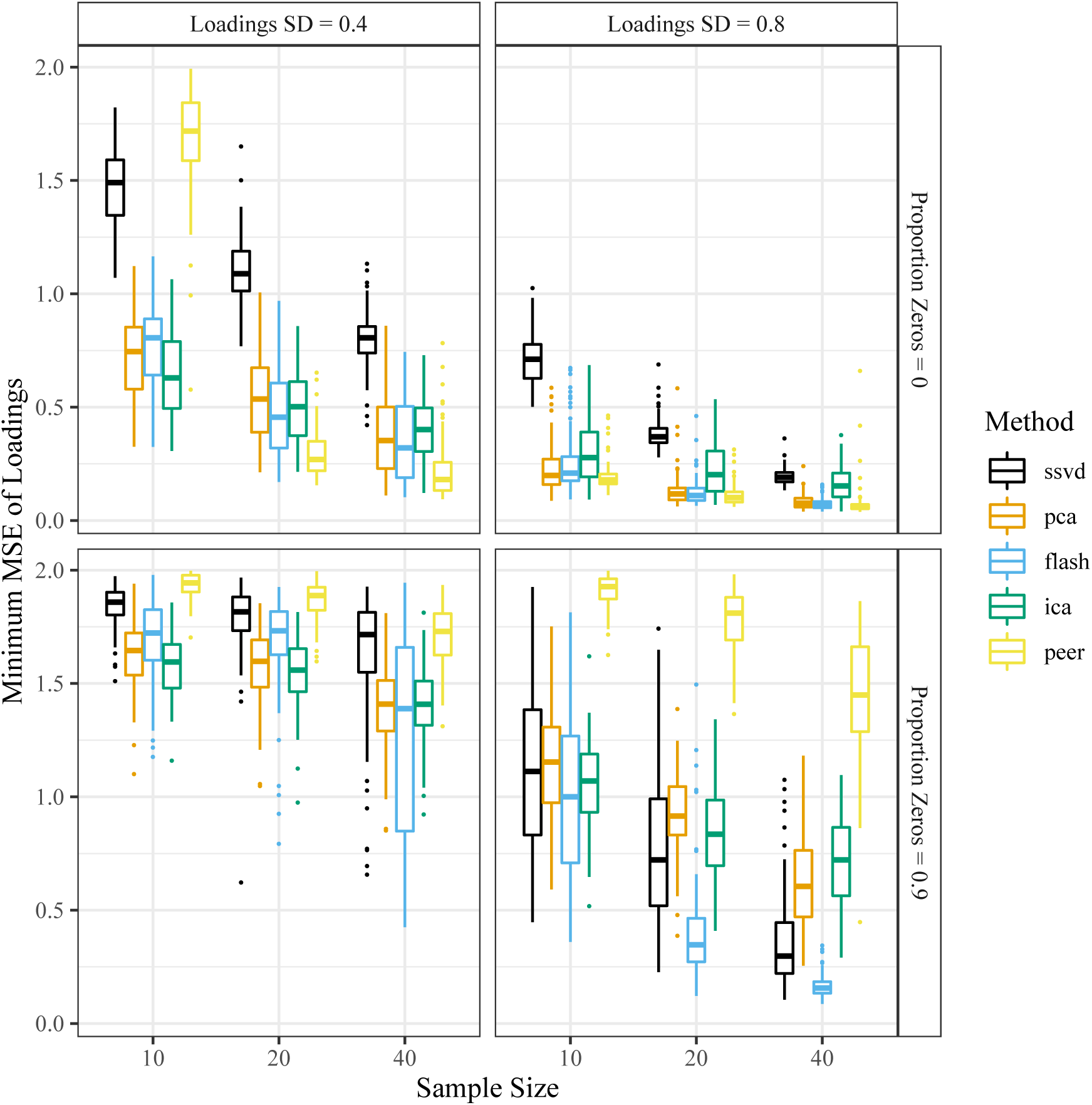
Minimum mean squared error between the added loading and the estimated loadings (*y*-axis), stratified by the sample size (*x*-axis), the factor analysis method (color), signal strength (column facets), and sparsity of the loadings (row facets). Before calculating the MSE, all loadings were scaled to have *ℓ*^2^-norm of 1. The target correlation between the added factor and the first unobserved factor was set to 0. A smaller minimum MSE is better.

**Figure S14:**
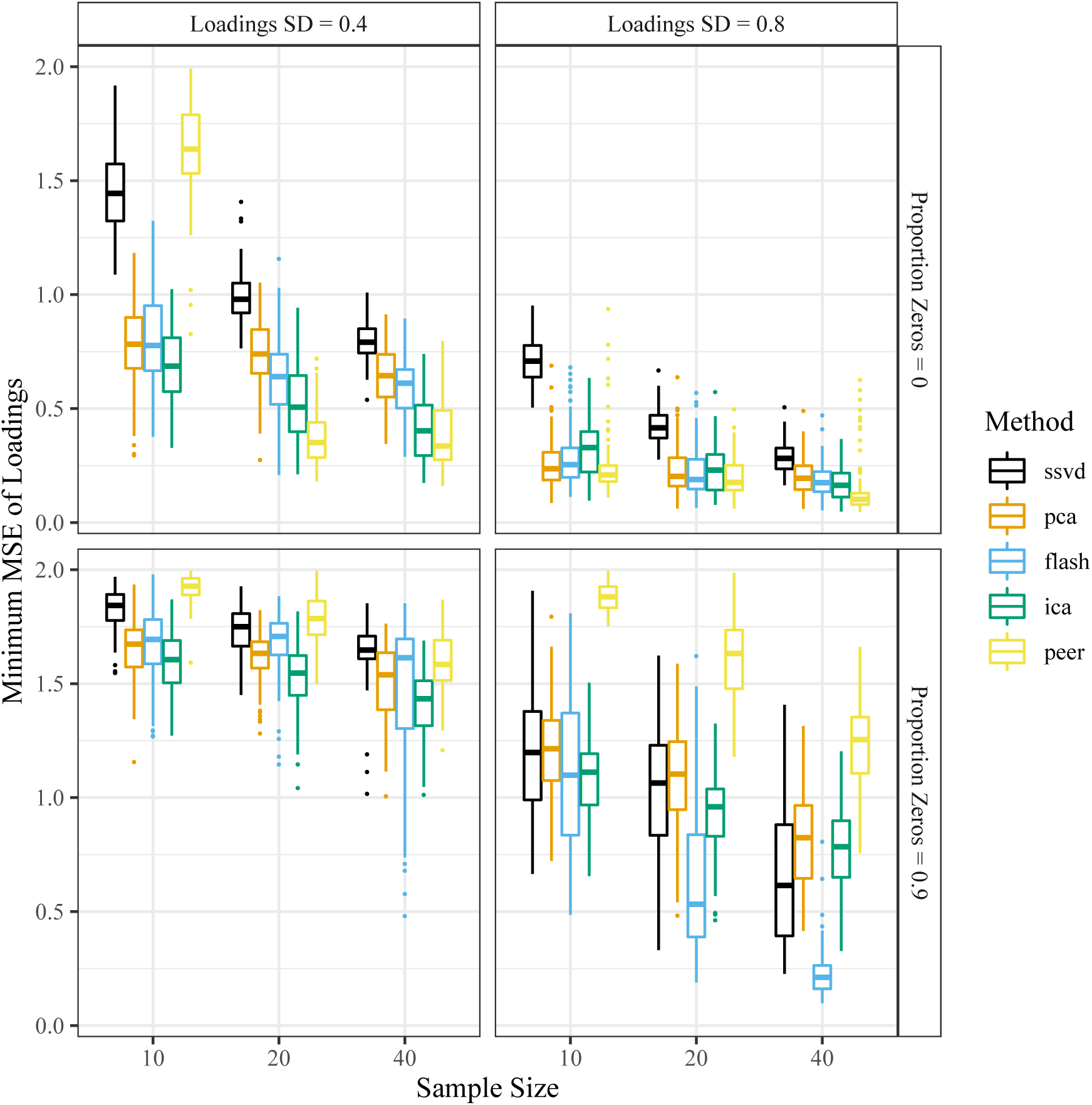
Minimum mean squared error between the added loading and the estimated loadings (*y*-axis), stratified by the sample size (*x*-axis), the factor analysis method (color), signal strength (column facets), and sparsity of the loadings (row facets). Before calculating the MSE, all loadings were scaled to have *ℓ*^2^-norm of 1. The target correlation between the added factor and the first unobserved factor was set to 0.5. A smaller minimum MSE is better.

**Figure S15:**
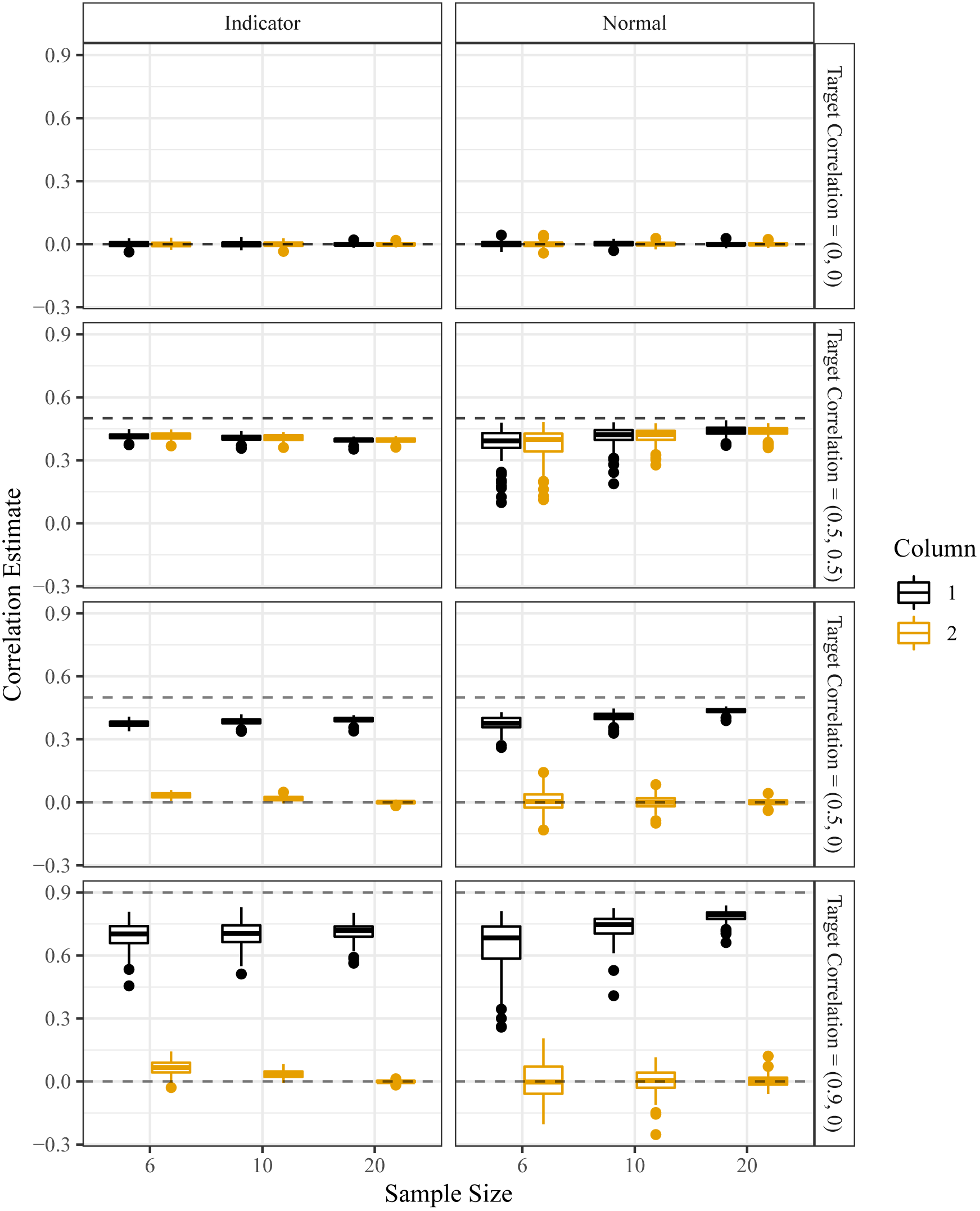
Boxplots of the Monte Carlo correlation estimator from Procedure 4 (*y*-axis), stratified by sample size (*x*-axis), the type of design matrix (column facets), the target correlations (row facets), and the column of the design matrix (color). Horizontal dashed lines are the two target correlations. The mean of the correlation estimates is approximately the true correlation.

